# From morphogenesis to pathogenesis: A cellulose loosening protein is one of the most widely distributed tools in nature

**DOI:** 10.1101/637728

**Authors:** William R. Chase, Olga Zhaxybayeva, Jorge Rocha, Daniel J. Cosgrove, Lori R. Shapiro

## Abstract

Plants must rearrange the network of complex carbohydrates in their cell walls during normal growth and development. To accomplish this, all plants depend on proteins called expansins that non-enzymatically loosen hydrogen bonds between cellulose microfibrils. Because of their key role in cell wall extension during growth, expansin genes are ubiquitous, diverse, and abundant throughout all land plants. Surprisingly, expansin genes have more recently been found in some bacteria and microbial eukaryotes, where their biological functions are largely unknown. Here, we reconstruct the phylogeny of microbial expansin genes. We find these genes in all eukaryotic microorganisms that have structural cellulose in their cell walls, suggesting expansins evolved in ancient marine microorganisms long before the evolution of land plants. We also find expansins in an unexpectedly high phylogenetic diversity of bacteria and fungi that do not have cellulosic cell walls. These bacteria and fungi with expansin genes inhabit varied ecological contexts mirroring the diversity of terrestrial and aquatic niches where plant and/or algal cellulosic cell walls are present. The microbial expansin phylogeny shows evidence of multiple horizontal gene transfer events within and between bacterial and eukaryotic microbial lineages, which may in part underlie their unusually broad phylogenetic distribution. Taken together, we find expansins to be unexpectedly widespread in both bacterial and eukaryotic genetic backgrounds, and that the contribution of these genes to bacterial and fungal ecological interactions with plants and algae has likely been underappreciated.

**Importance:** Cellulose is the most abundant biopolymer on earth. In plant cell walls, where most global cellulose biomass is found, cellulose microfibrils occur intertwined with hemicelluloses and pectins. The rigidity of this polysaccharide matrix provides plant cell walls with structural support, but this rigidity also restricts cellular growth and development. Irreversible, non-enzymatic loosening of structural carbohydrates by expansin proteins is key to successful cell wall growth in plants and green algae. Here, we find that expansin genes are distributed far more broadly throughout diverse bacterial and fungal lineages lacking cellulosic cell walls than previously known. Multiple horizontal gene transfer events are in part responsible for their unusually wide phylogenetic distribution. Together, these results suggest that in addition to being the key evolutionary innovation by which eukaryotes remodel structural cellulose in their cell walls, expansins likely have remarkably broad and under-recognized utility for microbial species that interact with plant and algal structural cellulose in diverse ecological contexts.

## Introduction

Cellulose – a linear polysaccharide comprised of hundreds to thousands of D-glucose units – is the most abundant biopolymer on Earth. The vast majority of global cellulose biomass is present in plant cell walls, where cellulose microfibrils are interlinked with hemicelluloses and pectins to provide structural support (1–3). This carbohydrate and protein matrix allows plant cell walls to withstand high tensile stresses, which can reach as high as 1000 atmospheres during growth (4). The strength of the structural carbohydrates in the cell wall also creates a formidable physical barrier against pathogenic microorganisms (5). Cellulosic cell walls similar in structure to those in plants also occur in some algal and other microbial eukaryotic groups (6–8). Tunicates (Urochordata) are the only metazoan group known to use cellulose structurally, and are thought to have acquired cellulose synthase genes horizontally from bacteria (9–11).

All of these diverse organisms are confronted with the dilemma of how to loosen their cellulose-based matrix of structural carbohydrates in order to expand their cell walls during normal growth and development. In plants and some green algae, non-enzymatic proteins called expansins provide the most important structural cellulose loosening functions, and are most highly expressed during active growth in any tissue where cell wall extension is critical (12–20). Expansin proteins are tightly packed two-domain structures of 200-250 amino acids with a planar polysaccharide binding surface (Supplemental Figure 1). The N-terminal domain is related to family 45 glycoside hydrolases, but lacks lytic activity. The C-terminal domain is related to group 2 grass pollen allergens (2, 21–25).

Expansin genes are universally present in all species of land plants, and most plant genomes contain multiple expansin homologs (26–28). In vascular plants, expansins have diversified from a common ancestor into four distinct genetic subfamilies. Two of these subfamilies, α and ϐ expansins, have been empirically shown to cause irreversible cell wall extension (2, 3, 26, 29, 30). The gene sequences of the two remaining subfamilies, expansin-like families A and B (EXLA and EXLB, respectively), contain both canonical expansin domains but no EXLA or EXLB have yet been functionally characterized. The current working hypothesis for expansin mode of action is non-enzymatic disruption of hydrogen bonds at biomechanical hotspots between cellulose microfibrils, or between cellulose microfibrils and hemicellulose. Interruption of hydrogen bonds allows slippage of carbohydrate polymers (*ie*, ‘loosening’) at load bearing elements of the cell wall, and is distinct from the action of hydrolytic enzymes. This structural carbohydrate loosening causes water uptake and irreversible extension of the cell wall without compromising tensile strength (19, 30, 31). The ubiquity of expansins in land plants and some green algae, the phylogenetic diversity of expansins in vascular plants, and their essential role in cell wall growth underlies the hypotheses that expansins may have first evolved in green algae and then diversified in land plants, and that these genes were necessary for the evolutionary success and adaptive radiation of the Plantae lineage (32–36).

What has long remained unknown is the distribution and function of expansin genes in non-Plantae organisms – especially those that do not have cellulosic cell walls (32, 37–40). Fungal and bacterial genes predicted to have similar structure as ϐ-expansins were first identified once databases began storing large numbers of genomic sequences. These microbial expansin-like genes were assigned to a newly established EXLX subfamily (22, 23). The first bacterial expansin gene with the predicted two domain structure of a plant expansin to be identified was a single copy expansin gene (Exlx1) from the rhizosphere plant commensal bacterium, *Bacillus subtilis*. This gene, referred to by expansin naming convention as BsExlx1, is highly divergent from plant expansins at the amino acid sequence level, but contains a conserved aspartic acid in domain 1 that is crucial for cell wall extension, and linear aromatic residues in domain 2 that are essential for polysaccharide binding (Supplemental Figure 1) (23, 38).

*Bacillus subtilis*, like many species of bacteria, utilizes cellulose as part of an extracellular biofilm matrix, but does not use cellulose as a cell wall structural component (41, 42). Functional characterization found that BsExlx1 significantly increases the efficiency of ephiphytic maize root colonization by *B. subtilis*, despite showing 10 times less *in vitro* cell wall loosening activity than plant expansins (23). This suggests that the function of expansins in microbial backgrounds may be to facilitate colonization of hosts that produce cellulosic cell walls (43–48). Since this first characterization of an expansin gene in a bacterium, additional EXLX family expansin genes have been identified in phylogenetically diverse bacteria, fungi and some microbial eukaryotes (reviewed in (32, 33)). Like plant expansins, no microbial expansins have been documented to have enzymatic activity (38, 49–54). Microbial expansin evolutionary history, taxonomic distribution, mechanism(s) of action, and ecological function(s) remains enigmatic in non-Plantae genetic backgrounds, and there is currently no framework for predicting their functional roles (38, 40, 51, 55–57).

Here, we examine the phylogenetic distribution of microbial expansin genes, and infer their possible ecological roles through four steps. First, we searched for expansin genes in all non-Plantae records of GenBank, and reconstruct the phylogeny of microbial expansin homologs in the context of the broader tree of life. Second, we consider the distribution of expansin genes relative to the life history of microorganisms that have them. Third, we analyze the microbial expansin phylogeny for signals of horizontal gene transfer, and hypothesize how ecological factors may be driving transfer of this gene between distantly related microbial taxa. Finally, we examine ongoing evolution through fusions with carbohydrate active proteins. Together, these analyses indicate that microbial expansins are more widely utilized by microorganisms than previously recognized, and that their distribution is in part driven by an underestimated importance for mediating bacterial and fungal interactions with live and dead plant and algal matter.

## Results

### Phylogenetic distribution of expansin genes across the tree of life

We surveyed the NCBI *nr* database (accessed Jan 2017) and identified a total of 600 unique expansin sequences in 491 species, in addition to those known in green algae (Chloroplastida) and terrestrial plants (Embryophyta) (Table 1, Supplemental Table 1). These 491 species with expansin homologs are comprised of macroscopic and microscopic organisms widely distributed across the tree of life (Figure 1). In Archaeplastida, expansins are present in red algae (Rhodophyta), which use cellulose as their main cell wall structural carbohydrate, and in *Cyanophora paradoxa*, the sole publicly available Glaucophyta genome sequence (Supplemental Table 2) (58). Glaucophyta are a rare and largely uncharacterized Archaeplastid group that likely diverged prior to the split between the red (Rhodophyta), and green (Chloroplastida) algal lineages (58, 59).

**Figure 1.**
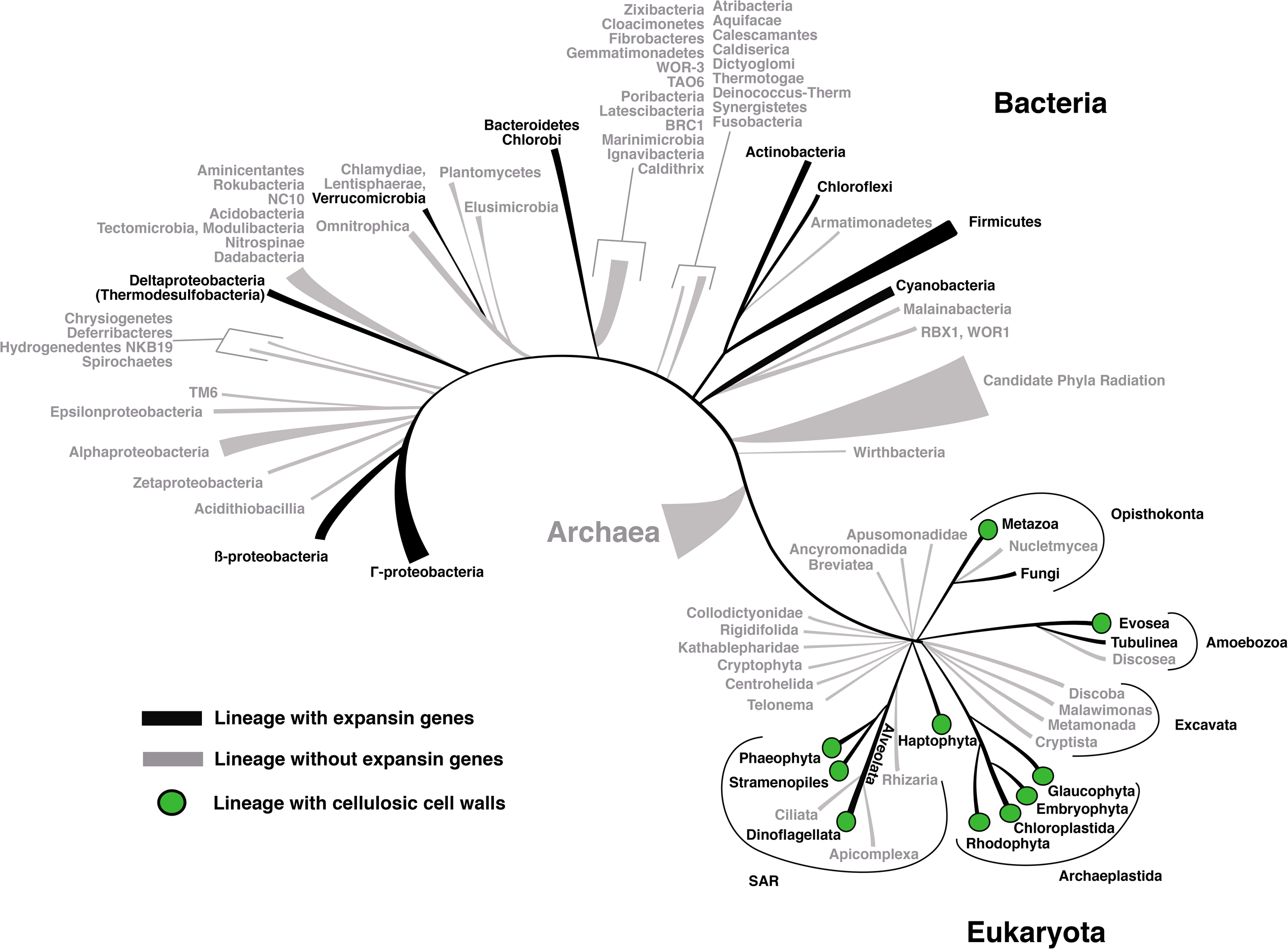
Distribution of expansins within major groups of the Tree of Life. The lineages within Eukaryota and Bacteria that have at least one species with an expansin gene are shown in black, and the lineages without a species with an expansin gene are in gray. No expansin homologs were detected in the available Archaeal genomes. Lineages with organisms that use cellulose structurally are marked with a green dot. The phyla relationships are based on (59, 123, 140) for Eukaryota and (124) for Bacteria.

Few expansin genes occur in Metazoans (Supplemental Table 3). Tunicates are the only metazoan group known to use cellulose structurally, and acquired their cellulose synthase genes horizontally from bacteria (11, 60, 61). *Oiklopeura dioica* is the sole tunicate species with a sequenced genome, and it contains an annotated expansin gene. Expansin genes are also annotated in several species of marine bivalves whose diets are partially plant matter or algae. For the sole glaucophyte and the few metazoans with annotated expansin homologs, it remains empirically unconfirmed whether their expansin genes are *bona fide* cellular genes, or sequencing contamination from digestive contents or other plant or microbial DNA (62, 63). Many plant pathogenic nematodes have proteins with partial functional and structural overlap with expansins, but their domain structure is reversed compared to the canonical plant and microbial expansin proteins. The evolutionary relationship of nematode expansin-like proteins to plant or microbial expansins remains unclear (30, 38, 64), and functional similarities in plant cell wall loosening function may be an example of convergent evolution.

In non-Archaeplastida eukaryotic microbes, Exlx homologs are present in both major lineages of Amoebozoa, one Alveolate (*Vitrella brassicaformis*), one Haptophyte (*Emiliania huxleyi*), and multiple lineages of Stramenopiles. Few species from these groups have sequenced and well-annotated genomes, and it is likely that more expansin homologs will be identified as more species from these lineages have their genomes sequenced and annotated. *E. huxleyi* and some Stramenopile lineages with expansin genes (such as Phaeophytes) are photosynthetic marine organisms with cellulosic cell walls. Many Amoebozoa and terrestrial Stramenopiles also have cellulosic cell walls. In the slime mold *Dictyostelium discoidum* (Amoebozoa), expansin genes are expressed while structural cellulose is being rearranged during fruiting body development (8). It is likely that *D. discoidum* – and other microbial eukaryotes with cellulosic cell walls – use expansins to modify their own structural cellulose. Many Oomycetes (a group of non-photosynthetic Stramenopiles) have expansin genes, but it remains unclear whether oomycetes use expansins for morphogenesis, interactions with plant cell walls, or both. All oomycetes with annotated expansin genes use cellulose structurally and have multiple expansin homologs per genome (Supplemental Table 1) which both suggest functions related to morphogenesis. However, these same oomycetes colonize plants as hosts and some are among the world’s worst agricultural plant pathogens, suggesting possible function(s) related to plant colonization (65–69).

Expansin genes are not detected in Archaea, but EXLX homologs are present in an unexpectedly diverse assortment of fungal and bacterial taxa. While many bacteria secrete cellulose as part of an extracellular biofilm matrix, none are known to utilize cellulose as a structural component of their cell walls (41, 42). Expansins are most abundant in Actinobacteria, Firmicutes, Myxobacteria, *γ*-Proteobacteria and ϐ-Proteobacteria, and are also present in some Chloroflexi, Bacteroidetes, Cyanobacteria and Verrucomicrobia. In fungi, none of which are known to use cellulose structurally, most expansin genes are in Ascomycota, and fewer expansins are in Basidiomycota, Chytrid fungi and symbiotic ectomycorrhizal fungi (Supplemental Table 4).

### Diversity of ecological niches inhabited by microbes with expansin genes

Since the first discovery of an expansin gene in a bacterium, it was hypothesized that most microbial expansins function as virulence factors (23, 25, 30, 33, 53). However, we find that only 28% (138 out of 491) of microbial species with expansins are plant pathogens (Figure 2, Supplemental Table 4). More than half (290 out of 491, or 59%) are described as plant commensals, soil inhabitants, or saprophytes – ecological contexts where microbes non-pathogenically interact with live plants and/or decaying plant matter (70). The remaining 13% of expansin-containing microbes inhabit a variety of terrestrial or aquatic ecological contexts, all of which have in common the presence of live or dead plant or algal matter.

**Figure 2.**
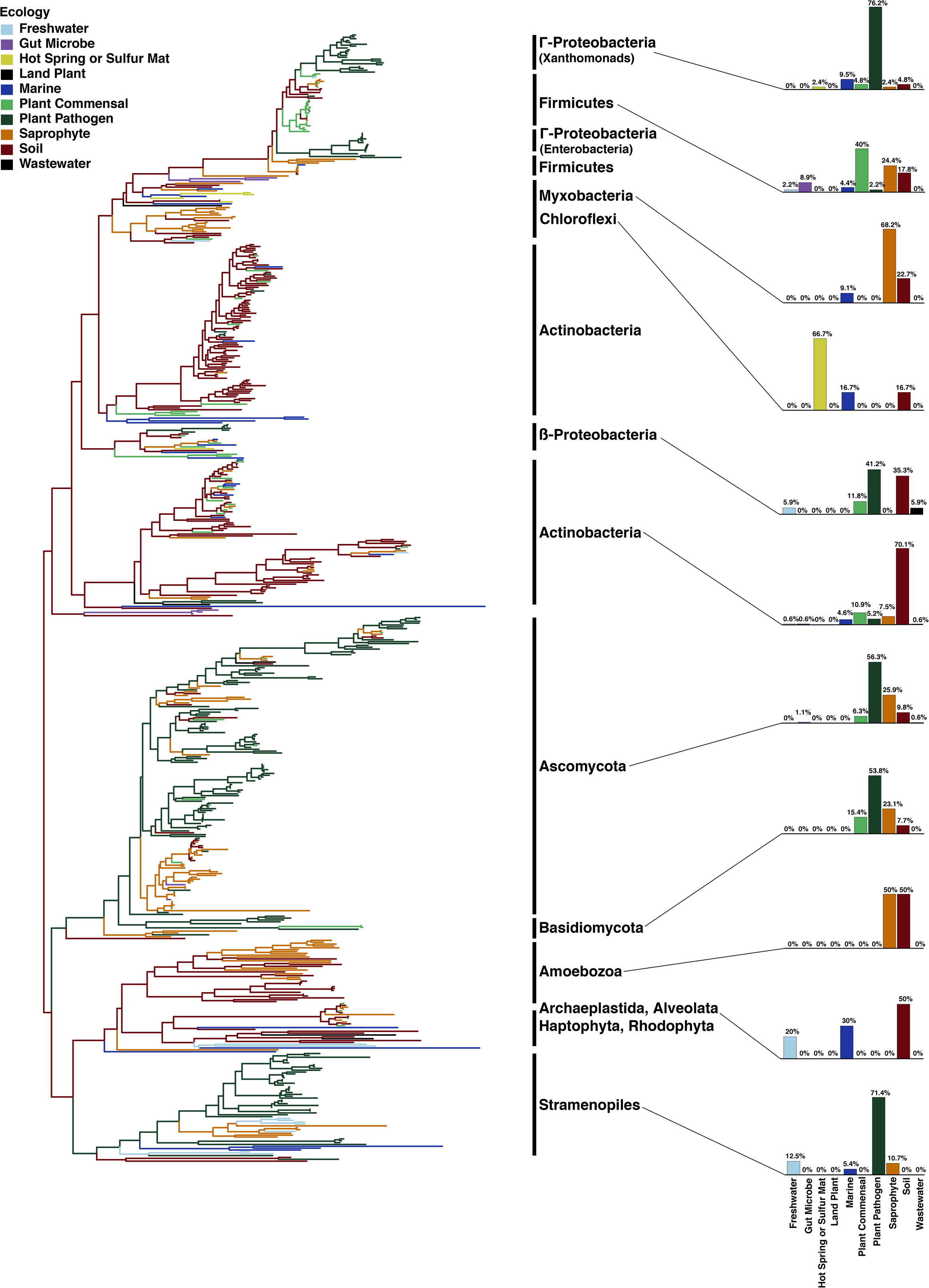
Ecological niches occupied by microbes with expansins. The maximum likelihood phylogenetic tree should be considered unrooted. Each branch is color-coded according to the ecological life history of that taxon (Supplementary Table 4). The major taxonomic groups of the tree are annotated with black bars. A proportional bar chart summarizing the distribution of ecological life histories is shown to the right of the major taxonomic groups. The scale bar, amino acid substitutions per site. See Supplemental Figure 2 for the tree with the individual taxa labels shown.

Bacteria comprise 61.3% (301 out of 491) of microbial species with expansin genes. Especially notable are Myxobacteria, where 85% (22 out of 26) of sequenced species have expansin genes (Table 1), suggesting that their ecological importance as saprophytes has been overshadowed by their use as laboratory models to understand bacterial multicellular behavior (71, 72). The Actinobacterial genera *Streptomyces, Nocardia*, and *Micromonospora* have many species with expansin genes. Numerous *Streptomyces* are plant growth promoters or pathogens, but *Nocardia* and *Micromonospora* are predominantly known as soil inhabitants and have few (or no) described plant associations (Supplemental Table 4) (73, 74). The high frequency of expansin genes suggests that microbial associations with live plants or dead plant matter is likely more common, and more ecologically important, for these actinobacterial genera than is currently recognized.

Many expansin genes are found in known plant growth promoting rhizobacteria, including strains of *Streptomyces, Bacillus, Micromonospora*, and *Rhizobacter*. In these species, expansin may function similarly to *B. subtilis* (23), and increase epiphytic colonization efficiency of plant roots. Only 15% of bacterial species (46 out of 301) with expansin genes are phytopathogens, and most of those pathogens occur in two *γ*-proteobacterial lineages, Xanthomonadaceae and Enterobacteriaceae. Other bacterial plant pathogens are sparsely scattered throughout the tree, and include economically important strains of *Ralstonia, Acidovorax, Streptomyces* and *Clavibacter michaganensis*. A conspicuous number of these expansin-containing bacteria are among the most economically costly agricultural plant pathogens (Table 2). Notably, all expansin-containing bacterial phytopathogens move systemically via xylem at some stage of pathogenesis – an unusual, highly virulent phenotype compared to localized lesions produced by most bacterial plant pathogens (75–82).

An additional 7% (22 out of 301) of bacterial species with expansins are marine or freshwater, and likely interact commensally with live algae or plant, or saprophytically degrade dead algal or plant matter. Several species of bacteria with expansin genes were isolated from plants growing in tidal flats, where they may facilitate plant-microbe symbiosis that allow both partners to better tolerate elevated salt levels (83). Expansins were also found from bacteria in acid mine drainage sites, sulfur mats and hot springs (84, 85). An expansin gene is present in *Cedecea neteri*, which has been isolated as both a plant commensal and a facultative termite gut symbiont (86–89). Expansin genes are found in bacteria (*Paenibacillus, Ruminococcus, Firmicutes, Actinobacteria*) and fungi (*Neocallimastix, Anaeromyces, Piromyces, Aspergillus*) that are commensals in herbivore ruminant guts and likely aid in degradation of ingested plant matter (Supplemental Table 4) (90).

Only 31.5% of microbes (155 out of 491) with an expansin homolog are fungi, and they inhabit a more restricted range of ecological habitats than bacteria. Almost all fungal species with expansin genes (94%; 146 out of 155) are described as plant pathogens, commensals, or saprophytes, although it is possible this reflects under-sampling of fungi compared to bacteria (91). A higher proportion of fungal species with expansins are phytopathogenic (52.3%; 81 out of 155) compared to the proportion of bacteria that are phytopathogens (15%; 46 out of 301). Expansin genes are present in many economically devastating fungal pathogens, including many that can cause vascular wilt diseases and can move via xylem during pathogenesis (Figure 2, Table 2, Supplemental Table 4) (92, 93).

### Horizontal gene transfer has shaped expansin distribution in microbes

Horizontal transfer of genes between distantly related organisms can introduce new traits and drive rapid evolutionary innovation in the recipient organism. In the microbial expansin phylogeny we identified 21 nodes that are in strong conflict with expected taxonomic relationships, which is suggestive of horizontal gene transfer. We then evaluated the statistical support for the relationships at these nodes with three phylogenetic analyses (Figure 3, Table 3). Some of these nodes are well-supported statistically, while others have low support in one or more tests, and verifying their placement will require better sampling and/or improved phylogenetic algorithms (Table 3).

**Figure 3.**
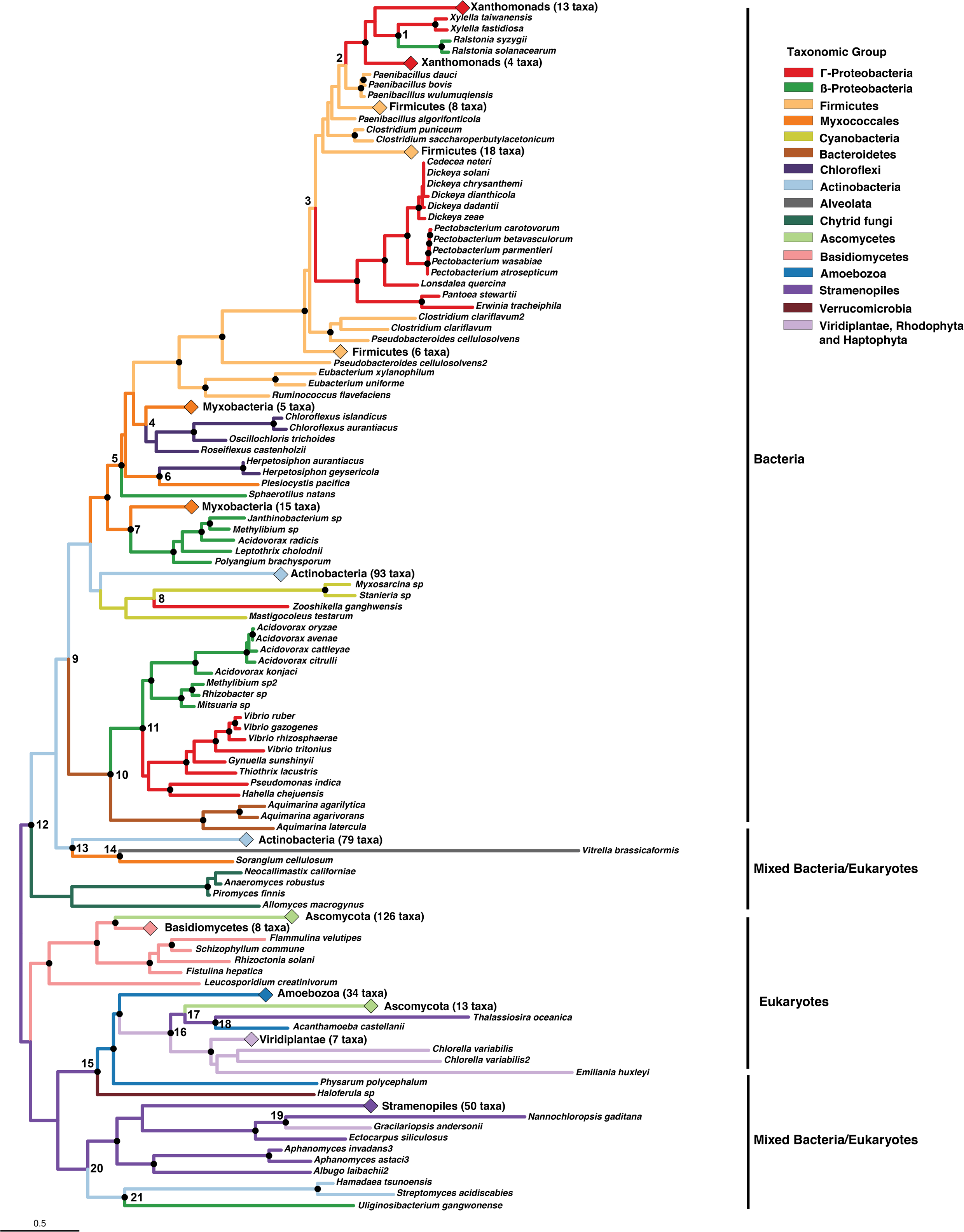
Evidence for horizontal exchange of expansin genes within and between Bacteria and Eukaryota. The maximum likelihood phylogenetic tree should be considered unrooted. Some groups are collapsed to improve presentation; these groups are marked with the number of taxa collapsed at that tip. Well-supported nodes (Shimodaira–Hasegawa like approximate likelihood ratio test > 70% and/or ultrafast bootstrap > 95%) are marked with black dots. Branches are colored according to taxonomic classification. Nodes inferred to be involved in the HGT events are shown in bold and numbered (1-21), which correspond to their entries in Table 3. Tree scale bar, number of amino acid substitutions per site for the expansin tree.

Four nodes represent putative intra-domain HGT events within Eukaryota (nodes 16, 17, 18, 19), twelve nodes represent putative intra-domain exchanges within Bacteria (nodes 1, 2, 3, 4, 5, 6, 7, 8, 9, 10, 11, 21), and five nodes represent putative inter-domain exchanges between Bacteria and Eukaryota (nodes 12, 13, 14, 15, 20). Within Eukaryota, the Rhodophyta red alga *Gracilariopsis andersonii* is recovered within Stramenopiles (node 19). Thirteen Ascomycota are recovered in a mixed group with Viridiplantae, the Amoebozoa *Acanthamoeba castellanii* and the Stramenopile *Thalassiosira oceanica* (nodes 16, 17, 18).

Of the 12 within-bacteria HGT events, five involve β-proteobacteria (nodes 1, 5, 7, 11, 21) and four involve *γ*-proteobacteria (nodes, 1, 2, 3, 8, 11). Two distinct groups of pathogenic *γ*-proteobacteria – the Xanthomonad and Enterobacterial plant pathogens – group with Firmicutes (nodes 2, 3). Similarities in ecological habitat and life histories between species at some nodes with putative HGT relationships – such as the marine *γ*-proteobacteria *Zooshikella ganghwensis* and marine Cyanobacteria at node 8, and plant pathogenic *Ralstonia* within plant pathogenic Xanthomonads at node 1 – suggest ecological niche may be a strong factor driving some expansin HGT events (94). Actinobacterial expansins separate into two main lineages, one comprised mainly of *Streptomyces* and the other mainly of *Micromonospora* and *Nocardia*.

These Actinobacterial lineages are separated by a polyphyletic group that includes β-proteobacteria, *γ*-proteobacteria and Bacteroidetes (nodes 9, 10, 11). The plant commensal *Acidovorax radicis* is part of the β-proteobacteria group recovered within Myxobacteria (node 7), while five other *Acidovorax* that are plant pathogens are in the β-proteobacteria group recovered in Actinobacteria (node 11). All Chloroflexi are recovered within Myxobacteria (nodes 4, 6).

Five of the 21 putative HGT events are inter-domain exchanges between Bacteria and Eukaryota. Node 15 recovers the expansin gene from *Haloferula* sp. (Verrucomicrobia), a marine symbiont of brown algae (Stramenopiles), near Amoebozoa and Stramenopiles (95). Node 12 groups Chytrid fungi as sister to Actinobacteria, and places the Chytrid expansins as basal to the bacterial expansins. The Alveolate *Vitrella brassicaformis* (node 14) is recovered as sister to an Actinobacteria-Myxobacteria intra-domain HGT event (node 13). Node 20 groups the expansin genes from two Actinobacteria (*Hamadaea tsuongensis* and *Streptomyces acidiscabies*) and a β-proteobacteria (*Uliginosibacterium gangwonense*) (node 21) with Stramenopiles. In previous studies, the *Streptomyces acidiscabies* expansin gene was recovered within a group of plant expansins, and this relationship was interpreted as phylogenetic evidence for an interdomain HGT from a green land plant donor to a bacterium (33). In our phylogeny – built with a much broader representation of Amoeboza and Stramenopiles than was available previously – this same *S. acidiscabies* expansin gene, plus an additional expansin gene from the actinobacterium *Hamadaea tsuonensis* – are still an example of interdomain HGT, but group with Stramenopiles and not within land plants. None of the 491 microbial expansin genes group within the Viridiplantae, strengthening the hypothesis that land plants were not the expansin gene donors to bacteria and fungi. Further, these five inter-domain HGT events (nodes 12, 13, 14, 15, 20) support the hypothesis that a bacterium could have acquired an expansin gene in a marine environment long before land plants evolved ~475-515 million years ago (96).

### Some microbial expansins co-occur with carbohydrate active proteins

In some fungi and bacteria, the two-domain canonical expansin gene is fused to additional glycoside hydrolase (GH) and/or carbohydrate binding module (CBMs). There are currently 83 recognized CBM families. All are non-enzymatic, and often function as part of a larger protein to facilitate adhesion to complex carbohydrates with high substrate specificity (24, 97). GHs are a group of enzymes widespread among plants and microbes that degrade complex carbohydrates, and are currently classified into 153 distinct families in the Carbohydrate Active Enzymes database (www.cazy.org). Out of 491 microbial species with expansin genes, 49 (9.9%) exist as fusions to a carbohydrate active domain (Figure 4, Supplemental Table 5). Fifteen of these fusions were previously known (33), and 34 are first identified here.

**Figure 4.**
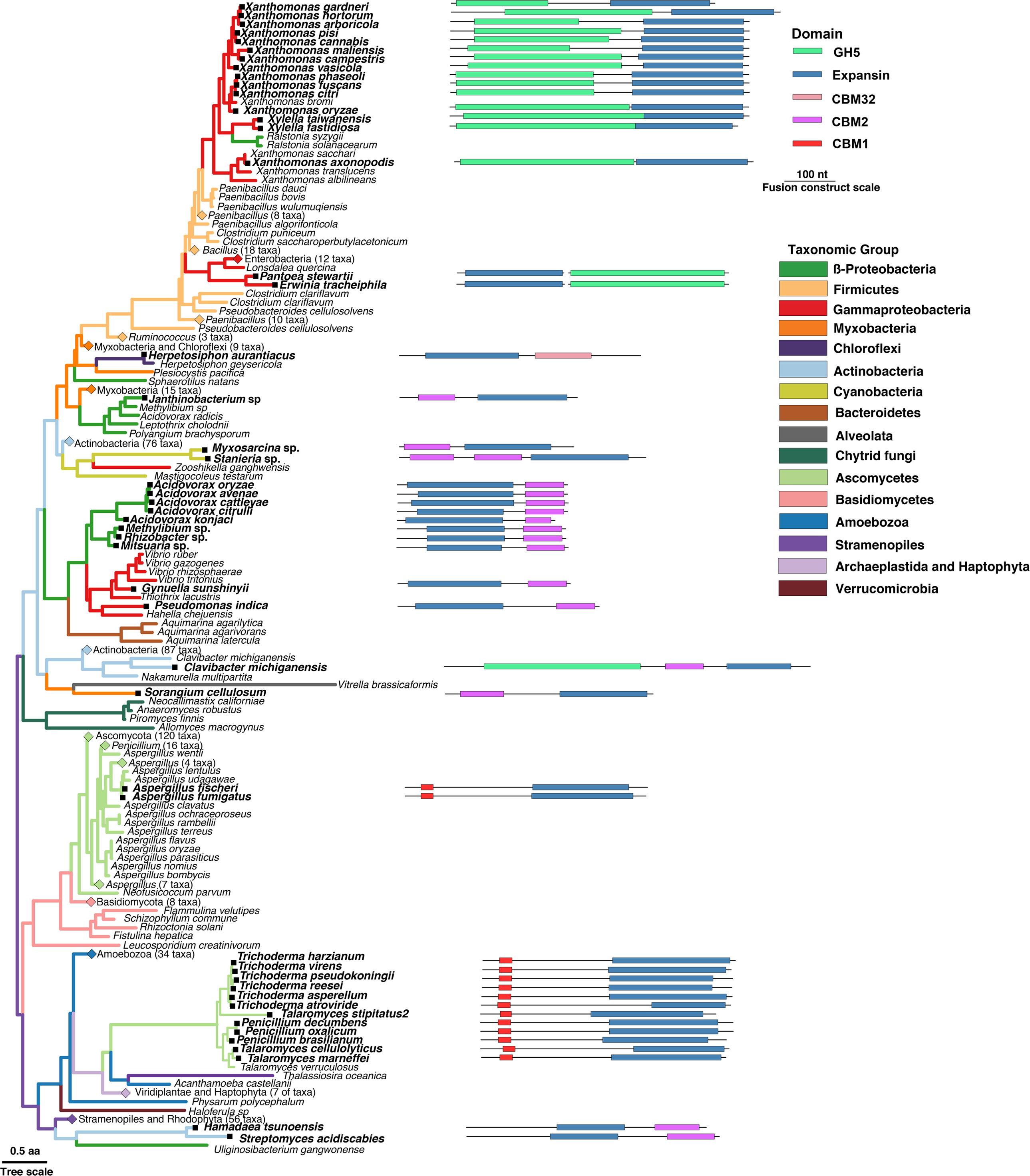
Gene architecture and phylogenetic distribution of expansin genes that are fused with carbohydrate active domains. Maximum likelihood phylogenetic tree of microbial expansin sequences should be considered unrooted. The tree branches are colored according to taxonomy. The tree scale bar is the number of amino acid substitutions per site. In 49 species where the expansin is fused to a carbohydrate active domain, the domain architectures are shown next to the taxa that have them. The domain architecture diagrams are drawn to scale, with the black line representing the length of the full nucleotide sequences of each gene, and carbohydrate active domains as colored rectangles. The domain architecture scale bar is the length (in nucleotides) of the expansin and carbohydrate active domains.

Carbohydrate binding module family 1 (CBM1) is the only carbohydrate active domain fused to fungal expansins (alternatively referred to in the literature as a ‘swollenins’ (32, 40, 55)). All 14 fungal species with expansin-CBM1 fusions are non-pathogenic. Twelve species of non-pathogenic *Trichoderma, Penicillium* and *Talaromyces* form a group distinct from the other predominantly pathogenic fungi without CBM1 fusions, and this group is a well-supported within-Eukaryota HGT event (Figure 3, node 17). Some *Trichoderma* spp., including those with expansin-CBM1 fusions, are among the most thoroughly characterized plant beneficial fungi (98, 99). We hypothesize that in fungal genetic backgrounds, expansin fusion to CBM1 increases fungal mutualistic capabilities to plant hosts, providing a selective advantage for fungal strains that contain this fusion.

In bacteria, expansins are predominantly found fused to domains from carbohydrate binding module family 2 (CBM2) and/or glycoside hydrolase family 5 (GH5). The Chloroflexi *Herpetosiphon aurantiacus* is the only microbe with an expansin fused to a CBM32 domain. A GH5-expansin fusion construct is present in 15 plant pathogenic Xanthomonadaceae. In some Cyanobacteria, β-proteobacteria, *γ*-proteobacteria and Actinobacteria, expansins are fused to CBM2 with variable domain arrangements (expansin-CBM2, CBM2-CBM2-expansin or CBM2-expansin). *Clavibacter michiganensis* is the only species with an expansin domain fused to both GH5 and CBM2 domains (GH5-CBM2-expansin domain arrangement). Most bacteria with unfused expansin genes are not plant pathogens (Figure 2, Supplemental Table 4). However, of the bacterial species with expansin fusions to GH5 and/or CBM2 domains, most (65.3%; 32 out of 49) are virulent phytopathogens (Supplemental Table 5). This suggests that in bacteria, expansin fusions are more likely than unfused expansins to function as a virulence factor.

The existence of variable fusion constructs (expansin-CBM32, GH5-expansin, expansin-CBM2, CBM2-expansin, CBM2-CBM2-expansin, and GH5-CBM2-expansin) indicates multiple independent origins of expansin fusions to carbohydrate active domains have occurred in bacteria and fungi. The repeated independent fusions of CBM2 and GH5 domains in bacteria, and only CBM1 in fungi – out of hundreds of CBM and GH families – suggests that CBM1, CBM2 and GH5 active domains in combination with expansin are uniquely useful for bacterial and fungal interactions with cellulosic cell walls.

The enterobacteria may offer mechanistic insight into how fusions can occur. In all enterobacteria, expansin genes are unfused to carbohydrate active domains. However, the plant pathogens *Erwinia tracheiphila* (80, 100) and *Pantoea stewartii* have a canonical expansin gene directly adjacent to – but in a separate open reading frame (ORF) – from a GH5 endoglucanase gene. This expansin-GH5 domain arrangement in *E. tracheiphila* and *P. stewartii* is in opposite positional order to the 15 Xanthomonadaceae with a GH5-expansin fusion construct, suggesting that in either *E. tracheiphila* or *P. stewartii* this gene architecture arose *de novo* and was not acquired horizontally from a Xanthomonadaceae donor. In *E. tracheiphila* and *P. stewartii*, both the expansin and GH5 ORFs have a predicted secretion signal peptide. The two coding sequences are separated by a stop codon and a short stretch of 40 nucleotides in *E. tracheiphila* and 51 in *P. stewartii* (KE136322.1, position 16101-17807 in *E. tracheiphila* (101) and NZ_CP017589.1, position 1851-3562 in *P. stewartii*). From this genetic architecture, it is possible that either a small deletion in the region between these ORFs, or a mutation in the stop codon separating them, could result in fusion of the two domains into a single gene (102).

## Discussion

We find that microbial expansin genes are more broadly distributed across diverse lineages of bacteria, fungi and other eukaryotic microbes than previously recognized. Especially notable is the presence of expansins in microbes inhabiting a previously unrecognized diversity of terrestrial and aquatic ecological niches, including those not traditionally thought of as cellulose-dominated. Many expansin genes are also found in microbes not yet known to interact with plants or algae, suggesting interactions with live or dead plant or algal matter is an overlooked yet important part of their ecological life histories. Identifying expansin genes in such a phylogenetically and ecologically diverse set of microbial species – including many which have not yet been described as interacting with plants or algae – suggests that the immense amount of global cellulose biomass (1) is an under-recognized selective pressure driving microbial evolution (94).

While the first organism to evolve an expansin gene and the timeframe of this innovation remains unknown, we hypothesize that the original expansin evolved long before the emergence of land plants 475-515 million years ago (96). Many microbes that use expansin proteins for cell wall expansion during growth and development – including Stramenopiles, Amoebozoa, Haptophyta, Alveolata, Rhodophyta and Chlorophyta – are lineages much older than land plants. The presence of expansin genes in all eukaryotic organisms with cellulosic cell walls, together with the absence of any extant alternate mechanism for irreversible cellulosic cell wall extension suggests that expansins may have been necessary for the success of the original organism with cellulosic cell walls (30, 96, 103, 104). This also raises the possibility that the EXLX microbial expansin subfamily could have been the first to evolve, and then diversified into distinct EXPA, EXPB, EXLA, and EXLB subfamilies in land plants. Ultimately, answering the question of expansin origin and ancient evolutionary dynamics will require greater taxon sampling, high confidence molecular dating of the different lineages with this gene, and accurate rooting of the expansin phylogeny.

The microbial expansin phylogeny indicates that HGT has been an important process shaping the distribution of expansins among microorganisms, and that expansin gene exchange is ongoing. In some cases, the presence of expansin genes in most sequenced species within a group (such as Myxobacteria, Xanthomonadaceae, and the *Pectobacterium* and *Dickeya* group of Entobacterial plant pathogens) suggests that original acquisition of an expansin likely occurred in a common ancestor of these taxa before these groups diversified (105). In other species – notably several plant pathogens – acquisition of an expansin likely occurred on more recent ecological time scales. In several bacterial pathogen species, acquisition of an expansin gene or gene fusion resulted in an ability to move systemically via xylem and achieve high within-host titre, which is a high virulence phenotype (51, 52, 77, 106–108). The high frequency of expansin genes in many virulent fungal and bacterial plant pathogens suggests that expansins or expansin fusions can function as a potent virulence factor when acquired by bacteria and fungi in simplified agro-ecosystems. The increase in virulence conferred by horizontal acquisition of expansins or expansin fusions by microbes in agricultural systems may amount to yet another demonstration of human-driven evolution of pathogenic micro-organisms (80, 109–111). The amenability of expansin genes to horizontal transfer between phylogenetically divergent microbial lineages, the functionality of this gene in diverse genetic backgrounds, and its repeated occurrence in virulent agricultural pathogens should elicit concern about the possibility of this gene facilitating the emergence of novel, highly virulent pathogen species or strains in managed agricultural settings.

In addition to expansin dissemination via HGT, functional evolution of microbial expansins is likely also driven by fusions with carbohydrate active proteins. There is a correlation between fusion to carbohydrate active domains with a transition between pathogenicity and commensalism, but in opposite directions for bacteria and fungi. In fungi, there was likely only one fusion of an expansin to a CBM1 domain. In bacteria, expansin genes have likely fused multiple times independently with a CBM2 domain, fused at least once to a GH5 domain in Xanthomonadaceae, and appears to be in a possible intermediate arrangement that may result in an additional fusion in *Erwinia tracheiphila* and/or *Pantoea stewartii*. The occurrence of expansin fusion constructs across the expansin phylogeny, repeated occurrence of fusions with the same carbohydrate active domains from multiple independent fusion events and the distinct ecological interactions of species with expansin fusions compared to closely related species with unfused expansin genes suggests that these fusion constructs have emergent (but still unknown) properties beyond their individual constituent domains.

We now recognize that all Eukaryotic microbes and macrobes in marine environments – and later in evolutionary history, on land – have evolved as part of complex multi-species communities (112–115). A mechanism to non-destructively manipulate structural cell wall cellulose would have been highly adaptive for the organism that first evolved a cellulosic cell wall. This same mechanism could have also been adaptive for the diverse microbes that colonize the surfaces of eukaryotes that have cellulosic cell walls as hosts. The functional flexibility of expansins – which are essential for normal growth and development in some lineages (land plants, red and green algae and some eukaryotic microbes) and accessory in others (bacteria, fungi and possibly other eukaryotic microbes) – appears unique. A more complete understanding of expansin evolutionary origin, functional diversification and emergent properties from fusions with carbohydrate active domains may offer unique insight into the origin of cellulosic cell walls, and mechanisms underlying host-microbe ecological interactions.

## Methods

### Plant and microbial expansin protein structures

The crystal structures of the bacterial expansin from *Bacillus subtilis* (BsEXLX1, PDB: 3D30) alone (116) and in complex with plant cellohexose (PDB: 4FER)(117), and the plant ϐ-expansin ZmEXPB1 (PDB: 2HCZ) from *Zea mays* (118) were downloaded from the Protein Data Bank (PDB) (119). The 3D protein structures were visualized with UCSF Chimera (v1.2.2) (120).

### Detection of microbial expansin sequences

Amino acid sequences encoding microbial expansins were identified in a two-step approach. In the first step, the NCBI non-redundant protein sequence database was queried using the keywords ‘expansin’ and ‘swollenin’ and excluding hits from Viridiplantae taxa (accessed Jan. 2017). The retrieved amino acid sequences were then curated to remove duplicates and ensure that all hits were *bona fide* microbial expansin genes. All hits were evaluated based on presence of the canonical expansin domains and key amino acid motifs (conserved aspartic acid in domain 1 and conserved aromatic triplet in domain 2) to the experimentally validated reference expansin sequences BsEXLX1 from *Bacillus subtilis* (AAB84448.1), and the alpha expansin AtEXPA4 from *Arabidopsis* (AEC09708.1). BsEXLX1 and AtEXPA4 were used as references because they represent the microbial expansin (BsEXLX1) or plant expansin (AtEXPA4) superfamilies, and the expansin function of both genes has been experimentally validated.

Records were removed from the dataset if the amino acid sequence lacked either of the characteristic expansin domains or key residues. Amino acids sequences that flanked the two canonical expansin domains that were included in the CDS record because of mis-annotation (such as RNA polymerase sequences) were trimmed so that only the canonical expansin domains (and carbohydrate associated domains, when present) were retained. Domains were annotated and identified using CD-Search from NCBI (30).

The expansin sequences retrieved in this initial, keyword-based search were then manually separated into bacteria, fungi, or microbial eukaryotic subsets. Representative sequences from these three taxonomic groups were then used as BLASTP queries to identify any microbial expansin gene sequences in the NCBI non-redundant (*nr*) database that may have been missed in the keyword search. The bacterial, fungal and eukaryotic microbe expansin sequences were used as BLASTP queries against the non-redundant *nr* protein database using default parameters, but excluding hits from Viridiplantae (121). This sequence-based approach yielded additional hits that were added to the existing sequence lists. The sequences were again aligned to the reference expansin sequences using MAFFT, and then manually filtered and trimmed to remove false positive hits and mis-annotated flanking regions. The final microbial expansin gene set contains 600 unique, *bona-fide* non-Viridiplantae expansin proteins from 491 distinct microbial species (Supplemental File 1). For 113 microbial species, there were at least two, and up to eight, non-identical expansin genes within the same species (Supplemental Table 1). All sequence alignments were performed using MAFFT (v. 7) with the options FFT-NS-i (122).

Taxonomy information was retrieved for all 491 microbial species. The presence of expansin genes was mapped onto the currently understood phylogenies for Eukaryota and Bacteria (59, 123–125). For each bacterial order with multiple species that contain expansin genes, the NCBI taxonomy database was used to determine the total number of named species and the total number of sequenced species (125).

### Phylogenetic tree reconstruction

Because of high amino acid divergence, the expansin homologs from bacteria, fungi and the other eukaryotic microbes were aligned separately using MAFFT (option E-INS-i) (126). Poorly aligned regions at the termini were manually trimmed to the point where a conserved block was shared across 90% of species. All three trimmed alignments were then combined, and aligned again with MAFFT (option E-INS-i). Viridiplantae expansin gene sequences from one dicot (AtEXPA4, GenBank: O48818.1), one monocot (ZmEXPB1, GenBank: P58738.2), one non-vascular plant (PpEXPA10, XP_024392378.1), four randomly chosen sequences from the charophyte green algae *Klebsormidium nitens* (GAQ91800.1, GAQ85527.1, GAQ79710.1, GAQ91109.1) and two sequences from the chlorophyte green algae *Chlorella variabilis* (XP_005846210.1, XP_005846208.1) were added to the dataset. The dataset was re-aligned a final time with MAFFT (option E-INS-i). The final sequence alignment contains 608 sequences and 689 amino acid sites (Supplemental File 1).

ModelFinder (as implemented within IQ-tree v. 1.6) was used to determine the best evolutionary model for the alignment (127). The WAG+R7 model was chosen as the best evolutionary model based on the Akaike information criterion (AIC). The phylogeny was reconstructed in IQ-tree (v. 1.6) (128), using a smaller perturbation strength and larger number of stop iterations (options -pers 0.2 -nstop 500) to avoid local minima (all other parameters default).

Node supports were estimated using the Shimodaira–Hasegawa like approximate likelihood ratio test (SH-aLRT) (129) and the ultrafast bootstrap with 1000 bootstrap pseudo-replicates (130). IQ-tree was run with these parameters 13 independent times to test the robustness of phylogenetic relationships. The resulting concensus tree of the 13 runs was manually rooted between prokaryotes and eukaryotes for presentation purposes (Supplemental File 2). The tree was visualized and annotated using the ggtree package in R (v. 3.4.2) (131).

### Inference of expansin horizontal gene transfer events

Twenty-one putative horizontal gene transfer events were identified by finding incongruences between the expansin gene tree (Supplemental Figure 2) and the species taxonomy. To further evaluate the strength of the relationships recovered at these 21 nodes, a Bayesian approach was used to reconstruct the microbial expansin gene tree. This was followed by pruning 259 taxa from the dataset, leaving 350 taxa in the reduced alignment (Supplemental File 3, Supplemental File 4), and rerunning a maximum likelihood analysis on the reduced dataset in IQ-tree with the same options and run parameters as the full tree (described above) (127). Using ModelFinder implemented within IQ-tree v. 1.6 (127), the WAG+G4 model was selected for the pruned alignment. For the Bayesian phylogeny, the expansins from *Vitrella brassicaformis* and *Emiliania huxleyi* were removed due to their extremely divergent sequences which may have interfered with convergence of the Bayesian model. MrBayes (v 3.2.6) (132) was then used to construct a Bayesian tree on the XSEDE cluster (133). Two independent runs were performed for 10 million generations, each with six chains using metropolis coupling with a heating parameter of 0.005 and swap frequency of 1. Each chain was sampled every 500 generations and the first 1.5 million samples were discarded as burn-in. The log likelihood of both runs plateaued after ~1.5 million generations (Supplemental Figure 6) and both runs converged on a similar tree (standard deviation of split frequencies between runs = 0.020738. All parameters of the MCMC algorithm are listed in Supplemental File 5. Because both runs converged on a similar tree, a majority rule consensus tree was constructed from the sampled trees of run 1 (Supplemental File 6).

### Ecological niche determination and phylogenetic tree annotation

For each microbial species with an expansin gene, a literature search was carried out to determine the known ecological associations (Supplemental Table 4). For many species, there was little or no documentation of the life history. Further, for many species, the existing descriptions of ecological life history may be incomplete. For example, we recognize that the classifications of ‘plant commensal’, ‘saprophyte’, and ‘soil dweller’ likely share significant functional overlap, and many microbes may fit multiple of these overlapping categories. Many microbes thought of (and researched) as ‘soil dwellers’ are likely also saprophytes, plant commensals, and/or plant pathogens depending on the environmental conditions (92, 93, 134, 135). Despite these caveats, each species was assigned to only one of the following ecological categories after evaluating the available ecological information: freshwater, marine, gut microbe, soil dweller, plant commensal, plant pathogen, saprophyte, hot spring, sulfur mat, or wastewater. The certainty (or lack thereof) for the ecological assignments for each species is also noted in Supplemental Table 4. The expansin microbial phylogeny was annotated with the collected ecological information in Supplemental Table 4 using the ggtree package in R (v. 3.4.2) (136).

### Identification of carbohydrate active domains fused to microbial expansin domains

A comprehensive list of microbial expansin genes fused to carbohydrate active domains was compiled by asearch of the *nr* database (121) with the keywords ‘expansin’ and ‘swollenin’. This was followed by a BLASTP search with the expansin-swollenin fusion from *Trichoderma reesei* (Accession number: CAB92328.1) as a query. Both searches were constrained to records with a bit score above 100 and more than 300 amino acid residues in length to exclude non-fused expansin genes, which arenormally ~200-250 amino acids in length. The matches that met these two criteria were retained as putative genes with expansin-carbohydrate fusions.

The presence of a carbohydrate active domain(s) was then evaluated with a batch CD-search (137) and dbsCAN search which identified any carbohydrate active domains(138). Records that shared more than 95% sequence identity to another record in the same species were considered redundant and were removed. The expansin domains were then aligned in MAFFT with the plant (AtEXPA4) and bacteria (BsEXLX1) reference expansin sequences (126). The expansin - carbohydrate active domain fusion constructs were plotted next to the expansin gene tree using ggtree and genoPlotR (139).

## Author Contributions

LRS conceived of the study, LRS and WRC designed analysis, WRC and LRS performed analyses; WRC, OZ, DJC, JR, LRS critically interpreted analyses, LRS wrote the first draft of the manuscript; WRC, OZ, DJC, JR, LRS contributed manuscript revisions.

## Data Deposition Statement

No new data were generated for this study. The data gathered from NCBI, and the custom scripts created for this study to analyze these data, are hosted at https://github.com/will-r-chase/microbe_expansin.

## Acknowledgements

OZ was supported in part by a Simons Foundation Young Investigator Award. JR was supported by Fundación Mexico en Harvard and Conacyt grant 237414; LRS was supported by NSF postdoctoral fellowship DBI-1202736. This project used the Extreme Science and Engineering Discovery Environment (XSEDE), which is supported by National Science Foundation grant number ACI-1548562. We thank Kerry E. Mauck and Rob Dunn for critical reading and constructive comments on the manuscript.

---

**Table 1. Distribution of microbial expansins in major taxonomic groups.**

**Table 2. Presence of expansin genes in the ‘Top 10’ most important species of plant pathogenic bacteria and fungi in a poll of plant pathologists by the journal Molecular Plant Pathology** (78, 141).

**Table 3. Maximum likelihood and Bayesian support values testing statistical strength at 21 nodes representing putative HGT events.**

## Supplemental Figure Legends

**Supplemental figure 1. Expansin protein structure.** (A) Surface of the bacterial expansin BsEXLX1 (PDB ID: 3D30). (B) Surface of the plant expansin ZmEXPB1 (PDB ID: 2HCZ). (C) Ribbon representation of the bacterial expansin BsEXLX1 in complex with cellohexaose (PDB ID: 4FER). On all panels, domains 1 and 2 are shown in dark grey and white, respectively; residues crucial for binding cellulose are colored in blue, while residues important for loosening of a cell wall are colored in yellow. On panel C, cellohexaose is shown in magenta.

**Supplemental figure 2. Phylogenetic relationship among all 600 microbial expansins inferred using maximum likelihood method.** The tree should be considered unrooted. Each branch is color-coded according to the taxonomic affiliation of the organism. Well-supported nodes of the tree (Shimodaira–Hasegawa like approximate likelihood ratio test > 70% or ultrafast bootstrap > 95%) are marked with black dots. The nodes where relationships conflict with expected taxonomic relationships are numbered as in Figure 3 and Table 3. The scale bar, amino acid substitutions per site.

**Supplemental figure 3. Phylogenetic relationships among the detected microbial expansins inferred using Bayesian approach.** The tree is the majority rule consensus of the trees obtained via Bayesian inference. Numbers at the nodes represent posterior probabilities of the nodes. The tree should be considered unrooted. Each branch is color-coded according to the taxonomic affiliation of an organism. The scale bar, amino acid substitutions per site.

**Supplemental figure 4. Phylogenetic relationships among the 350 selected microbial expansins inferred using maximum likelihood method.** The tree should be considered unrooted. Each branch is color-coded according to the taxonomic affiliation of an organism. Support values for each node are from Shimodaira–Hasegawa like approximate likelihood ratio test and ultrafast bootstrap analysis. The scale bar, amino acid substitutions per site.

**Supplemental figure 5: NCBI Common Tree representation of the species tree for all 491 microbial species with expansin genes**

**Supplemental figure 6. Trace plot of two independent runs of Bayesian inference.** Two runs reached stationary phase after ~1.5 million generations.

**Supplemental table 1: List of microbial species with multiple expansin genes per genome.**

**Supplemental table 2. Number of publicly available genomes in the eukaryotic groups depicted in Figure 1.**

**Supplemental table 3. Animals whose genomes are known to contain at least one expansin gene.**

**Supplemental table 4. Ecological metadata for each microbial species with an expansin gene.**

**Supplemental table 5. Metadata for the 49 expansins that are fused with carbohydrate active domains.**

## References

1. Bar-On YM, Phillips R, Milo R. 2018. The biomass distribution on Earth. Proceedings of the National Academy of Sciences:201711842.

2. Cosgrove DJ. 2000. Loosening of plant cell walls by expansins. Nature 407:321–326.

3. Li Y, Jones L, McQueen-Mason S. 2003. Expansins and cell growth. Current Opinion in Plant Biology 6:603–610.

4. Cosgrove DJ. 2005. Growth of the plant cell wall. Nature Reviews 6:850–861.

5. Hématy K, Cherk C, Somerville S. 2009. Host–pathogen warfare at the plant cell wall. Current Opinion in Plant Biology 12:406–413.

6. Popper ZA, Michel G, Hervé C, Domozych DS, Willats WG, Tuohy MG, Kloareg B, Stengel DB. 2011. Evolution and diversity of plant cell walls: from algae to flowering plants. Annual Review of Plant Biology 62:567–90.

7. Popper ZA, Tuohy MG. 2010. Beyond the green: understanding the evolutionary puzzle of plant and algal cell walls. Plant Physiology 153:373–383.

8. Darley CP, Li Y, Schaap P, McQueen-Mason SJ. 2003. Expression of a family of expansin - like proteins during the development of *Dictyostelium discoideum*. FEBS Letters 546:416–418.

9. Kimura S, Itoh T. 1995. Evidence for the role of the glomerulocyte in cellulose synthesis in the tunicate, M*etandrocarpa uedai*. Protoplasma 186:24–33.

10. Tamai N, Tatsumi D, Matsumoto T. 2004. Rheological properties and molecular structure of tunicate cellulose in LiCl/1, 3-dimethyl-2-imidazolidinone. Biomacromolecules 5:422–432.

11. Nakashima K, Yamada L, Satou Y, Azuma J-i, Satoh N. 2004. The evolutionary origin of animal cellulose synthase. Development Genes and Evolution 214:81–88.

12. Cho H-T, Cosgrove DJ. 2000. Altered expression of expansin modulates leaf growth and pedicel abscission in *Arabidopsis thaliana*. Proceedings of the National Academy of Sciences 97:9783–9788.

13. Pien S, Wyrzykowska J, McQueen-Mason S, Smart C, Fleming A. 2001. Local expression of expansin induces the entire process of leaf development and modifies leaf shape. Proceedings of the National Academy of Sciences 98:11812–11817.

14. Lee D-K, Ahn JH, Song S-K, Do Choi Y, Lee JS. 2003. Expression of an expansin gene is correlated with root elongation in soybean. Plant Physiology 131:985–997.

15. Gray-Mitsumune M, Mellerowicz EJ, Abe H, Schrader J, Winzéll A, Sterky F, Blomqvist K, McQueen-Mason S, Teeri TT, Sundberg B. 2004. Expansins abundant in secondary xylem belong to subgroup A of the α-expansin gene family. Plant Physiology 135:1552–1564.

16. Im K-H, Cosgrove DJ, Jones AM. 2000. Subcellular localization of expansin mRNA in xylem cells. Plant Physiology 123:463–470.

17. Brummell DA, Harpster MH, Civello PM, Palys JM, Bennett AB, Dunsmuir P. 1999. Modification of expansin protein abundance in tomato fruit alters softening and cell wall polymer metabolism during ripening. The Plant Cell 11:2203–2216.

18. Chen F, Bradford KJ. 2000. Expression of an expansin is associated with endosperm weakening during tomato seed germination. Plant Physiology 124:1265–1274.

19. Cosgrove DJ. 2016. Catalysts of plant cell wall loosening. F1000Research 5.

20. Ramakrishna P, Duarte PR, Rance GA, Schubert M, Vordermaier V, Dai Vu L, Murphy E, Barro AV, Swarup K, Moirangthem K. 2019. EXPANSIN A1-mediated radial swelling of pericycle cells positions anticlinal cell divisions during lateral root initiation. Proceedings of the National Academy of Sciences:201820882.

21. Yennawar NH, Li L-C, Dudzinski DM, Tabuchi A, Cosgrove DJ. 2006. Crystal structure and activities of EXPB1 (Zea m 1), a β-expansin and group-1 pollen allergen from maize. Proceedings of the National Academy of Sciences 103:14664–14671.

22. Kende H, Bradford K, Brummell D, Cho H-T, Cosgrove D, Fleming A, Gehring C, Lee Y, McQueen-Mason S, Rose J. 2004. Nomenclature for members of the expansin superfamily of genes and proteins. Plant Molecular Biology 55:311–314.

23. Kerff F, Amoroso A, Herman R, Sauvage E, Petrella S, Filée P, Charlier P, Joris B, Tabuchi A, Nikolaidis N, Cosgrove DJ. 2008. Crystal structure and activity of *Bacillus subtilis* YoaJ (EXLX1), a bacterial expansin that promotes root colonization. Proceedings of the National Academy of Sciences 105:16876–16881.

24. Lombard V, Golaconda Ramulu H, Drula E, Coutinho PM, Henrissat B. 2013. The carbohydrate-active enzymes database (CAZy) in 2013. Nucleic Acids Research 42:D490–D495.

25. Sampedro J, Cosgrove DJ. 2005. The expansin superfamily. Genome biology 6:242.

26. Li Y, Darley CP, Ongaro V, Fleming A, Schipper O, Baldauf SL, McQueen-Mason SJ. 2002. Plant expansins are a complex multigene family with an ancient evolutionary origin. Plant Physiology 128:854–864.

27. Ding A, Marowa P, Kong Y. 2016. Genome-wide identification of the expansin gene family in tobacco (*Nicotiana tabacum*). Molecular Genetics and Genomics 291:1891–1907.

28. Santiago TR, Pereira VM, de Souza WR, Steindorff AS, Cunha BA, Gaspar M, Fávaro LC, Formighieri EF, Kobayashi AK, Molinari HB. 2018. Genome-wide identification, characterization and expression profile analysis of expansins gene family in sugarcane (*Saccharum* spp.). PloS One 13:e0191081.

29. Lee Y, Choi D, Kende H. 2001. Expansins: ever-expanding numbers and functions. Current Opinion in Plant Biology 4:527–532.

30. Cosgrove DJ. 2015. Plant expansins: diversity and interactions with plant cell walls. Current Opinion in Plant Biology 25:162–172.

31. Wang T, Park YB, Caporini MA, Rosay M, Zhong L, Cosgrove DJ, Hong M. 2013. Sensitivity-enhanced solid-state NMR detection of expansin’s target in plant cell walls. Proceedings of the National Academy of Sciences doi: 10.1073/pnas.1316290110:201316290.

32. Cosgrove DJ. 2017. Microbial Expansins. Annual Review of Microbiology 71:479–497.

33. Nikolaidis N, Doran N, Cosgrove DJ. 2013. Plant expansins in bacteria and fungi: evolution by horizontal gene transfer and independent domain fusion. Molecular Biology and Evolution 31:376–386.

34. Carey RE, Hepler NK, Cosgrove DJ. 2013. *Selaginella moellendorffii* has a reduced and highly conserved expansin superfamily with genes more closely related to angiosperms than to bryophytes. BMC Plant Biology 13:4.

35. Schipper O, Schaefer D, Reski R, Fleming A. 2002. Expansins in the bryophyte *Physcomitrella patens*. Plant Molecular Biology 50:789–802.

36. Carey RE, Cosgrove DJ. 2007. Portrait of the expansin superfamily in *Physcomitrella patens:* comparisons with angiosperm expansins. Annals of Botany 99:1131–1141.

37. Nikolaidis N, Doran N, Cosgrove DJ. 2014. Plant expansins in bacteria and fungi: evolution by horizontal gene transfer and independent domain fusion. Molecular biology and evolution 31:376–386.

38. Georgelis N, Nikolaidis N, Cosgrove DJ. 2015. Bacterial expansins and related proteins from the world of microbes. Applied Microbiology and Biotechnology 99:3807–3823.

39. Ogasawara S, Shimada N, Kawata T. 2009. Role of an expansin - like molecule in *Dictyostelium morphogenesis* and regulation of its gene expression by the signal transducer and activator of transcription protein Dd - STATa. Development, Growth & Differentiation 51:109–122.

40. Brotman Y, Briff E, Viterbo A, Chet I. 2008. Role of swollenin, an expansin-like protein from *Trichoderma*, in plant root colonization. Plant Physiology 147:779–789.

41. Branda SS, Vik Å, Friedman L, Kolter R. 2005. Biofilms: the matrix revisited. Trends in microbiology 13:20–26.

42. Schleifer KH, Kandler O. 1972. Peptidoglycan types of bacterial cell walls and their taxonomic implications. Bacteriological Reviews 36:407.

43. Burke C, Thomas T, Lewis M, Steinberg P, Kjelleberg S. 2011. Composition, uniqueness and variability of the epiphytic bacterial community of the green alga *Ulva australis*. The ISME Journal 5:590.

44. Egan S, Harder T, Burke C, Steinberg P, Kjelleberg S, Thomas T. 2013. The seaweed holobiont: understanding seaweed–bacteria interactions. FEMS Microbiology Reviews 37:462–476.

45. Haney CH, Samuel BS, Bush J, Ausubel FM. 2015. Associations with rhizosphere bacteria can confer an adaptive advantage to plants. Nature Plants 1.

46. Niu B, Paulson JN, Zheng X, Kolter R. 2017. Simplified and representative bacterial community of maize roots. Proceedings of the National Academy of Sciences:201616148.

47. Shabat SKB, Sasson G, Doron-Faigenboim A, Durman T, Yaacoby S, Miller MEB, White BA, Shterzer N, Mizrahi I. 2016. Specific microbiome-dependent mechanisms underlie the energy harvest efficiency of ruminants. The ISME Journal 10:2958.

48. Meibom KL, Li XB, Nielsen AT, Wu CY, Roseman S, Schoolnik GK. 2004. The *Vibrio cholerae* chitin utilization program. Proceedings of the National Academy of Sciences 101:2524.

49. Cosgrove DJ. 2017. Microbial Expansins. Annual Review of Microbiology 71.

50. Tancos MA, Lowe-Power TM, Peritore-Galve FC, Tran TM, Allen C, Smart CD. 2017. Plant - like bacterial expansins play contrasting roles in two tomato vascular pathogens. Molecular Plant Pathology.

51. Jahr H, Dreier J, Meletzus D, Bahro R, Eichenlaub R. 2000. The endo-β-1, 4-glucanase CelA of *Clavibacter michiganensis* subsp. *michiganensis* is a pathogenicity determinant required for induction of bacterial wilt of tomato. Molecular Plant-Microbe Interactions 13:703–714.

52. Olarte-Lozano M, Mendoza-Nuñez MA, Pastor N, Segovia L, Folch-Mallol J, Martínez-Anaya C. 2014. PcExl1 a novel acid expansin-like protein from the plant pathogen *Pectobacterium carotovorum*, binds cell walls differently to BsEXLX1. PLoS One 9:e95638.

53. Laine MJ, Haapalainen M, Wahlroos T, Kankare K, Nissinen R, Kassuwi S, Metzler MC. 2000. The cellulase encoded by the native plasmid of *Clavibacter michiganensis* ssp. *sepedonicus* plays a role in virulence and contains an expansin-like domain. Physiological and Molecular Plant Pathology 57:221–233.

54. Tancos MA, Lowe-Power TM, Peritore-Galve FC, Tran TM, Allen C, Smart CD. 2018. Plant-like bacterial expansins play contrasting roles in two tomato vascular pathogens. Molecular Plant Pathology 19:1210–1221.

55. Saloheimo M, Paloheimo M, Hakola S, Pere J, Swanson B, Nyyssönen E, Bhatia A, Ward M, Penttilä M. 2002. Swollenin, a *Trichoderma reesei* protein with sequence similarity to the plant expansins, exhibits disruption activity on cellulosic materials. European Journal of Biochemistry 269:4202–4211.

56. Hwang IS, Oh E-J, Lee HB, Oh C-S. 2018. Functional characterization of two cellulase genes in the Gram-positive pathogenic bacterium *Clavibacter michiganensis* for wilting in tomato. Molecular Plant-Microbe Interactions.

57. Junior AT, Dolce LG, de Oliveira Neto M, Polikarpov I. 2015. *Xanthomonas campestris* expansin-like X domain is a structurally disordered beta-sheet macromolecule capable of synergistically enhancing enzymatic efficiency of cellulose hydrolysis. Biotechnology Letters 37:2419–2426.

58. Price DC, Chan CX, Yoon HS, Yang EC, Qiu H, Weber AP, Schwacke R, Gross J, Blouin NA, Lane C. 2012. *Cyanophora paradoxa* genome elucidates origin of photosynthesis in algae and plants. Science 335:843–847.

59. Adl SM, Bass D, Lane CE, Lukeš J, Schoch CL, Smirnov A, Agatha S, Berney C, Brown MW, Burki F. 2019. Revisions to the classification, nomenclature, and diversity of eukaryotes. Journal of Eukaryotic Microbiology 66:4–119.

60. Sasakura Y, Ogura Y, Treen N, Yokomori R, Park S-J, Nakai K, Saiga H, Sakuma T, Yamamoto T, Fujiwara S. 2016. Transcriptional regulation of a horizontally transferred gene from bacterium to chordate. Proceedings of the Royal Society B: Biological Sciences 283:20161712.

61. Sasakura Y, Nakashima K, Awazu S, Matsuoka T, Nakayama A, Azuma J-i, Satoh N. 2005. Transposon-mediated insertional mutagenesis revealed the functions of animal cellulose synthase in the ascidian *Ciona intestinalis*. Proceedings of the National Academy of Sciences 102:15134–15139.

62. Coutteau P, Sorgeloos P. 1992. The use of algal substitutes and the requirement for live algae in the hatchery and nursery rearing of bivalve molluscs: an international survey. Journal of Shellfish Research.

63. Sakamoto K, Touhata K, Yamashita M, Kasai A, Toyohara H. 2007. Cellulose digestion by common Japanese freshwater clam *Corbicula japonica*. Fisheries Science 73:675–683.

64. Danchin EG, Rosso M-N, Vieira P, de Almeida-Engler J, Coutinho PM, Henrissat B, Abad P. 2010. Multiple lateral gene transfers and duplications have promoted plant parasitism ability in nematodes. Proceedings of the National Academy of Sciences 107:17651–17656.

65. Fawke S, Doumane M, Schornack S. 2015. Oomycete interactions with plants: infection strategies and resistance principles. Microbiology and Molecular Biology Reviews 79:263–280.

66. Kamoun S. 2001. Nonhost resistance to *Phytophthora:* novel prospects for a classical problem. Current Opinion in Plant Biology 4:295–300.

67. Hardham AR. 2007. Cell biology of plant–oomycete interactions. Cellular Microbiology 9:31–39.

68. Grenville-Briggs LJ, Anderson VL, Fugelstad J, Avrova AO, Bouzenzana J, Williams A, Wawra S, Whisson SC, Birch PR, Bulone V. 2008. Cellulose synthesis in *Phytophthora infestans* is required for normal appressorium formation and successful infection of potato. The Plant Cell 20:720–738.

69. Helbert W, Sugiyama J, Ishihara M, Yamanaka S. 1997. Characterization of native crystalline cellulose in the cell walls of Oomycota. Journal of Biotechnology 57:29–37.

70. Lofgren LA, LeBlanc NR, Certano AK, Nachtigall J, LaBine KM, Riddle J, Broz K, Dong Y, Bethan B, Kafer CW. 2018. Fusarium graminearum: pathogen or endophyte of North American grasses? New Phytologist.

71. Crespi BJ. 2001. The evolution of social behavior in microorganisms. Trends in Ecology & Evolution 16:178–183.

72. Neil RB, Hite D, Kelrick MI, Lockhart ML, Lee K. 2005. Myxobacterial biodiversity in an established oak-hickory forest and a savanna restoration site. Current Microbiology 50:88–95.

73. Loria R, Kers J, Joshi M. 2006. Evolution of plant pathogenicity in *Streptomyces*. Annual Review of Phytopathology 44:469–487.

74. Goodfellow M, Williams S. 1983. Ecology of Actinomycetes. Annual Reviews in Microbiology 37:189–216.

75. Agrios. 2005. Plant pathology, http://doc18.rupdfbook.com/plant-pathology-fifth-edition-by-george-n-agrios-PDFs-179089.pdf.

76. Bae C, Han SW, Song Y-R, Kim B-Y, Lee H-J, Lee J-M, Yeam I, Heu S, Oh C-S. 2015. Infection processes of xylem-colonizing pathogenic bacteria: possible explanations for the scarcity of qualitative disease resistance genes against them in crops. Theoretical and Applied Genetics 128:1219–1229.

77. Ewald PW. 1993. Evolution of infectious disease. Oxford University Press.

78. Mansfield J, Genin S, Magori S, Citovsky V, Sriariyanum M, Ronald P, Dow M, Verdier V, Beer SV, Machado MA. 2012. Top 10 plant pathogenic bacteria in molecular plant pathology. Molecular Plant Pathology 13:614–629.

79. Shapiro L, De Moraes CM, Stephenson AG, Mescher MC. 2012. Pathogen effects on vegetative and floral odours mediate vector attraction and host exposure in a complex pathosystem. Ecology Letters 15:1430–1438.

80. Shapiro LR, Paulson JN, Arnold BJ, Scully ED, Zhaxybayeva O, Pierce N, Rocha J, Klepac-Ceraj V, Holton K, Kolter R. 2018. An introduced crop plant is driving diversification of the virulent bacterial pathogen *Erwinia tracheiphila*. mBio 9:e01307–18.

81. Roper MC. 2011. *Pantoea stewartii* subsp. *stewartii:* lessons learned from a xylem-dwelling pathogen of sweet corn. Molecular Plant Pathology 12:628–637.

82. Smith EF. 1920. An introduction to bacterial diseases of plants. W.B. Saunders Company, Philadelphia.

83. Marasco R, Rolli E, Ettoumi B, Vigani G, Mapelli F, Borin S, Abou-Hadid AF, El-Behairy UA, Sorlini C, Cherif A. 2012. A drought resistance-promoting microbiome is selected by root system under desert farming. PLoS One 7:e48479.

84. Sgroy V, Cassán F, Masciarelli O, Del Papa MF, Lagares A, Luna V. 2009. Isolation and characterization of endophytic plant growth-promoting (PGPB) or stress homeostasis-regulating (PSHB) bacteria associated to the halophyte *Prosopis strombulifera*. Applied Microbiology and Biotechnology 85:371–381.

85. Chung EJ, Park JA, Jeon CO, Chung YR. 2015. *Gynuella sunshinyii* gen. nov., sp. nov., an antifungal rhizobacterium isolated from a halophyte, *Carex scabrifolia* Steud. International Journal of Systematic and Evolutionary Microbiology 65:1038–1043.

86. Thong-On A, Suzuki K, Noda S, Inoue J-i, Kajiwara S, Ohkuma M. 2012. Isolation and characterization of anaerobic bacteria for symbiotic recycling of uric acid nitrogen in the gut of various termites. Microbes and Environments 27:186–192.

87. Chan K-G, Tan K-H, Yin W-F, Tan J-Y. 2014. Complete genome sequence of *Cedecea neteri* strain SSMD04, a bacterium isolated from pickled mackerel sashimi. genomeA 2:e01339–14.

88. Farmer J, Sheth NK, Hudzinski JA, Rose HD, Asbury MF. 1982. Bacteremia due to *Cedecea neteri* sp. nov. Journal of Clinical Microbiology 16:775–778.

89. Aguilera A, Pascual J, Loza E, Lopez J, Garcia G, Liaño F, Quereda C, Ortuño J. 1995. Bacteraemia with *Cedecea neteri* in a patient with systemic lupus erythematosus. Postgraduate Medical Journal 71:179.

90. Morrison M, Pope PB, Denman SE, McSweeney CS. 2009. Plant biomass degradation by gut microbiomes: more of the same or something new? Current Opinion in Biotechnology 20:358–363.

91. Amend A, Burgaud G, Cunliffe M, Edgcomb VP, Ettinger CL, Gutiérrez MH, Heitman J, Hom EFY, Ianiri G, Jones AC, Kagami M, Picard KT, Quandt CA, Raghukumar S, Riquelme M, Stajich J, Vargas-Muñiz J, Walker AK, Yarden O, Gladfelter AS. 2019. Fungi in the marine environment: open questions and unsolved problems. mBio 10:e01189–18.

92. Lofgren LA, LeBlanc NR, Certano AK, Nachtigall J, LaBine KM, Riddle J, Broz K, Dong Y, Bethan B, Kafer CW. 2018. *Fusarium graminearum:* pathogen or endophyte of North American grasses? New Phytologist 217:1203–1212.

93. Selosse MA, Schneider-Maunoury L, Martos F. 2018. Time to re - think fungal ecology? Fungal ecological niches are often prejudged. New Phytologist 217:968–972.

94. Smillie CS, Smith MB, Friedman J, Cordero OX, David LA, Alm EJ. 2011. Ecology drives a global network of gene exchange connecting the human microbiome. Nature 480:241–244.

95. Mancuso FP, D’Hondt S, Willems A, Airoldi L, De Clerck O. 2016. Diversity and temporal dynamics of the epiphytic bacterial communities associated with the canopy-forming seaweed *Cystoseira compressa* (Esper) Gerloff and Nizamuddin. Frontiers in Microbiology 7.

96. Morris JL, Puttick MN, Clark JW, Edwards D, Kenrick P, Pressel S, Wellman CH, Yang Z, Schneider H, Donoghue PCJ. 2018. The timescale of early land plant evolution. Proceedings of the National Academy of Sciences 15:E2274–E2283.

97. Boraston AB, Bolam DN, Gilbert HJ, Davies GJ. 2004. Carbohydrate-binding modules: fine-tuning polysaccharide recognition. Biochemical Journal 382:769–781.

98. Harman GE, Howell CR, Viterbo A, Chet I, Lorito M. 2004. *Trichoderma* species— opportunistic, avirulent plant symbionts. Nature Reviews Microbiology 2:43.

99. Howell C. 2003. Mechanisms employed by *Trichoderma* species in the biological control of plant diseases: the history and evolution of current concepts. Plant Disease 87:4–10.

100. Shapiro LR, Scully ED, Straub TJ, Park J, Stephenson AG, Beattie GA, Gleason ML, Kolter R, Coelho MC, Moraes CMD, Mescher MC, Zhaxybayeva O. 2016. Horizontal gene acquisitions, mobile element proliferation, and genome decay in the host-restricted plant pathogen *Erwinia tracheiphila*. Genome Biology and Evolution 8:649–664.

101. Shapiro LR, Scully ED, Roberts D, Straub TJ, Geib SM, Park J, Stephenson AG, Rojas ES, Liu Q, Beattie G, Gleason M, Moraes CMD, Mescher MC, Fleischer SJ, Kolter R, Pierce N, Zhaxybayeva O. 2015. Draft genome sequence of *Erwinia tracheiphila*, an economically important bacterial pathogen of cucurbits. genomeA 3:e00482–15.

102. Kummerfeld SK, Teichmann SA. 2005. Relative rates of gene fusion and fission in multi-domain proteins. Trends in Genetics 21:25–30.

103. Vannerum K, Huysman MJ, De Rycke R, Vuylsteke M, Leliaert F, Pollier J, Lütz-Meindl U, Gillard J, De Veylder L, Goossens A. 2011. Transcriptional analysis of cell growth and morphogenesis in the unicellular green alga *Micrasterias* (Streptophyta), with emphasis on the role of expansin. BMC Plant Biology 11:128.

104. Van de Poel B, Cooper ED, Van Der Straeten D, Chang C, Delwiche CF. 2016. Transcriptome profiling of the green alga *Spirogyra pratensis* (Charophyta) suggests an ancestral role for ethylene in cell wall metabolism, photosynthesis, and abiotic stress responses. Plant Physiology 172:533–545.

105. McDonald BR, Currie CR. 2017. Lateral gene transfer dynamics in the ancient bacterial genus *Streptomyces*. mBio 8:e00644–17.

106. Ewald PW. 1994. Evolution of infectious disease. Oxford University Press on Demand.

107. Mennerat A, Nilsen F, Ebert D, Skorping A. 2010. Intensive farming: evolutionary implications for parasites and pathogens. Evolutionary Biology 37:59–67.

108. Pulkkinen K, Suomalainen L-R, Read A, Ebert D, Rintamäki P, Valtonen E. 2010. Intensive fish farming and the evolution of pathogen virulence: the case of columnaris disease in Finland. Proceedings of the Royal Society of London B: Biological Sciences 277:593–600.

109. Mira A, Pushker R, Rodríguez-Valera F. 2006. The Neolithic revolution of bacterial genomes. Trends in Microbiology 14:200–206.

110. Stukenbrock EH, Bataillon T. 2012. A population genomics perspective on the emergence and adaptation of new plant pathogens in agro-ecosystems. PLoS Pathogens 8:el002893.

111. McDonald BA, Stukenbrock EH. 2016. Rapid emergence of pathogens in agro-ecosystems: global threats to agricultural sustainability and food security. Philosophical transactions - Royal Society Biological sciences 371:20160026.

112. Segev E, Wyche TP, Kim KH, Petersen J, Ellebrandt C, Vlamakis H, Barteneva N, Paulson JN, Chai L, Clardy J, Kolter R. 2016. Dynamic metabolic exchange governs a marine algal-bacterial interaction. eLife 5:e17473.

113. Moran M, Belas R, Schell M, Gonzalez J, Sun F, Sun S, Binder B, Edmonds J, Ye W, Orcutt B. 2007. Ecological genomics of marine Roseobacters. Appied and Environmental Microbiology 73:4559–4569.

114. Kolter R, Chimileski S. 2019. The end of microbiology. Environmental Microbiology 20:1955–1959.

115. Chimileski S, Kolter R. 2017. Life at the Edge of Sight: A Photographic Exploration of the Microbial World. Harvard University Press.

116. Kerff F, Amoroso A, Herman R, Sauvage E, Petrella S, Filée P, Charlier P, Joris B, Tabuchi A, Nikolaidis N. 2008. PBD ID: 3D30 Crystal structure and activity of *Bacillus subtilis* YoaJ (EXLX1), a bacterial expansin that promotes root colonization. Proceedings of the National Academy of Sciences 105:16876–16881.

117. Georgelis N, Yennawar NH, Cosgrove DJ. 2012. (PDB: 4FER) Structural basis for entropy-driven cellulose binding by a type-A cellulose-binding module (CBM) and bacterial expansin. Proceedings of the National Academy of Sciences:201213200.

118. Yennawar NH, Li L-C, Dudzinski DM, Tabuchi A, Cosgrove DJ. 2006. (PDB: 2HCZ) Crystal structure and activities of EXPB1 (Zea m 1), a β-expansin and group-1 pollen allergen from maize. Proceedings of the National Academy of Sciences 103:14664–14671.

119. Rose PW, Prlić A, Altunkaya A, Bi C, Bradley AR, Christie CH, Costanzo LD, Duarte JM, Dutta S, Feng Z, Green RK, Goodsell DS, Hudson B, Kalro T, Lowe R, Peisach E, Randle C, Rose AS, Shao C, Tao Y-P, Valasatava Y, Voigt M, Westbrook JD, Woo J, Yang H, Young JY, Zardecki C, Berman HM, Burley. SK. 2017. The RCSB protein data bank: integrative view of protein, gene and 3D structural information. Nucleic Acids Research 45:D271–D281.

120. Pettersen EF, Goddard TD, Huang CC, Couch GS, Greenblatt DM, Meng EC, Ferrin TE. 2004. UCSF Chimera—a visualization system for exploratory research and analysis. Journal of Computational Chemistry 25:1605–1612.

121. Altschul SF, Gish W, Miller W, Myers EW, Lipman DJ. 1990. Basic local alignment search tool. Journal of Molecular Biology 215:403–410.

122. Katoh K, Standley DM. 2013. MAFFT multiple sequence alignment software version 7: improvements in performance and usability. Molecular Biology and Evolution 30:772–780.

123. Adl SM, Simpson AG, Lane CE, Lukeš J, Bass D, Bowser SS, Brown MW, Burki F, Dunthorn M, Hampl V. 2012. The revised classification of eukaryotes. Journal of Eukaryotic Microbiology 59:429–514.

124. Hug LA, Baker BJ, Anantharaman K, Brown CT, Probst AJ, Castelle CJ, Butterfield CN, Hernsdorf AW, Amano Y, Ise K. 2016. A new view of the tree of life. Nature Microbiology 1:16048.

125. Federhen S. 2011. The NCBI taxonomy database. Nucleic Acids Research 40:D136–D143.

126. Katoh K, Misawa K, Kuma Ki, Miyata T. 2002. MAFFT: a novel method for rapid multiple sequence alignment based on fast Fourier transform. Nucleic Acids Research 30:3059–3066.

127. Kalyaanamoorthy S, Minh BQ, Wong TK, von Haeseler A, Jermiin LS. 2017. ModelFinder: fast model selection for accurate phylogenetic estimates. Nature Methods 14:587.

128. Nguyen L-T, Schmidt HA, von Haeseler A, Minh BQ. 2014. IQ-TREE: a fast and effective stochastic algorithm for estimating maximum-likelihood phylogenies. Molecular Biology and Evolution 32:268–274.

129. Guindon S, Dufayard J-F, Lefort V, Anisimova M, Hordijk W, Gascuel O. 2010. New algorithms and methods to estimate maximum-likelihood phylogenies: assessing the performance of PhyML 3.0. Systematic Biology 59:307–321.

130. Hoang DT, Chernomor O, von Haeseler A, Minh BQ, Le SV. 2017. UFBoot2: Improving the Ultrafast Bootstrap Approximation. Molecular biology and evolution:msx281.

131. Team RC. 2015. R: A language and environment for statistical computing. R Foundation of Statistical Computing Vienna, Austria.

132. Ronquist F, Huelsenbeck JP. 2003. MRBAYES 3: Bayesian phylogenetic inference under mixed models. Bioinformatics 19:1572–1574.

133. Towns J, Cockerill T, Dahan M, Foster I, Gaither K, Grimshaw A, Hazlewood V, Lathrop S, Lifka D, Peterson GD, Roskies R, Scott JR, Wilkins-Diehr N. 2014. XSEDE: Accelerating Scientific Discovery. Computing in Science & Engineering 16:62–74.

134. Chen K-H. 2017. Evolution of fungal endophytes and their functional transitions between endophytism and saprotrophism.

135. Carroll G. 1988. Fungal endophytes in stems and leaves: from latent pathogen to mutualistic symbiont. Ecology 69:2–9.

136. Yu G, Smith DK, Zhu H, Guan Y, Lam TTY. 2017. ggtree: an R package for visualization and annotation of phylogenetic trees with their covariates and other associated data. Methods in Ecology and Evolution 8:28–36.

137. Marchler-Bauer A, Bryant SH. 2004. CD-Search: protein domain annotations on the fly. Nucleic Acids Research 32:W327–W331.

138. Yin Y, Mao X, Yang J, Chen X, Mao F, Xu Y. 2012. dbCAN: a web resource for automated carbohydrate-active enzyme annotation. Nucleic Acids Research 40:W445–W451.

139. Guy L, Kultima JR, Andersson SG. 2010. genoPlotR: comparative gene and genome visualization in R. Bioinformatics 26:2334–2335.

140. Pánek T, Zadrobílková E, Walker G, Brown MW, Gentekaki E, Hroudová M, Kang S, Roger AJ, Tice AK, Vlček Č. 2016. First multigene analysis of Archamoebae (Amoebozoa: Conosa) robustly reveals its phylogeny and shows that Entamoebidae represents a deep lineage of the group. Molecular Phylogenetics and Evolution 98:41–51.

141. Dean R, Van Kan JA, Pretorius ZA, Hammond-Kosack KE, Di Pietro A, Spanu PD, Rudd JJ, Dickman M, Kahmann R, Ellis J. 2012. The ‘Top 10’ fungal pathogens in molecular plant pathology. Molecular Plant Pathology 13:414–430.

