## Supplementary figures and images for "From morphogenesis to pathogenesis: A cellulose loosening protein is one of the most widely distributed tools in nature"

### Supplemental_Figure1

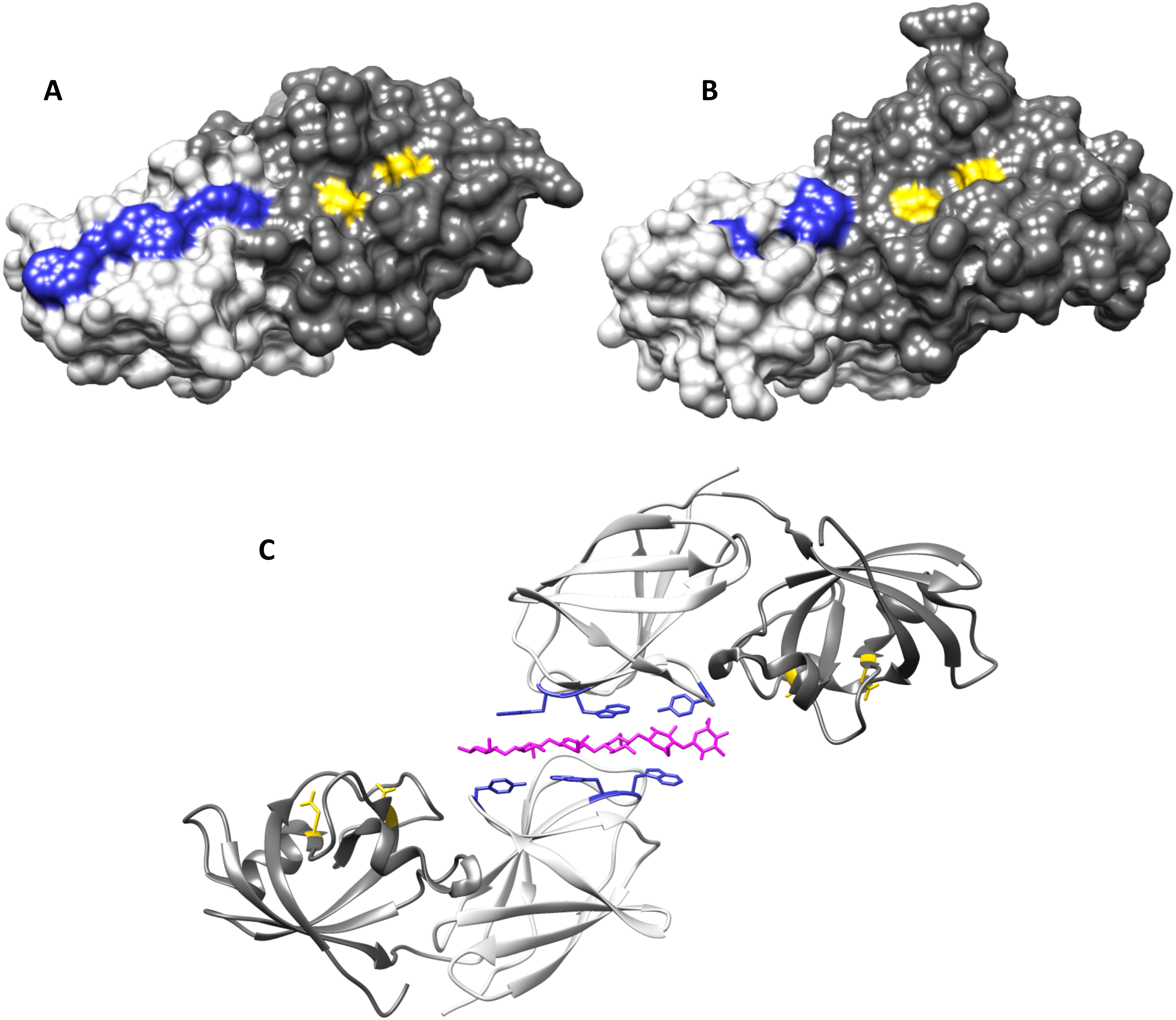

### Supplemental_Figure2

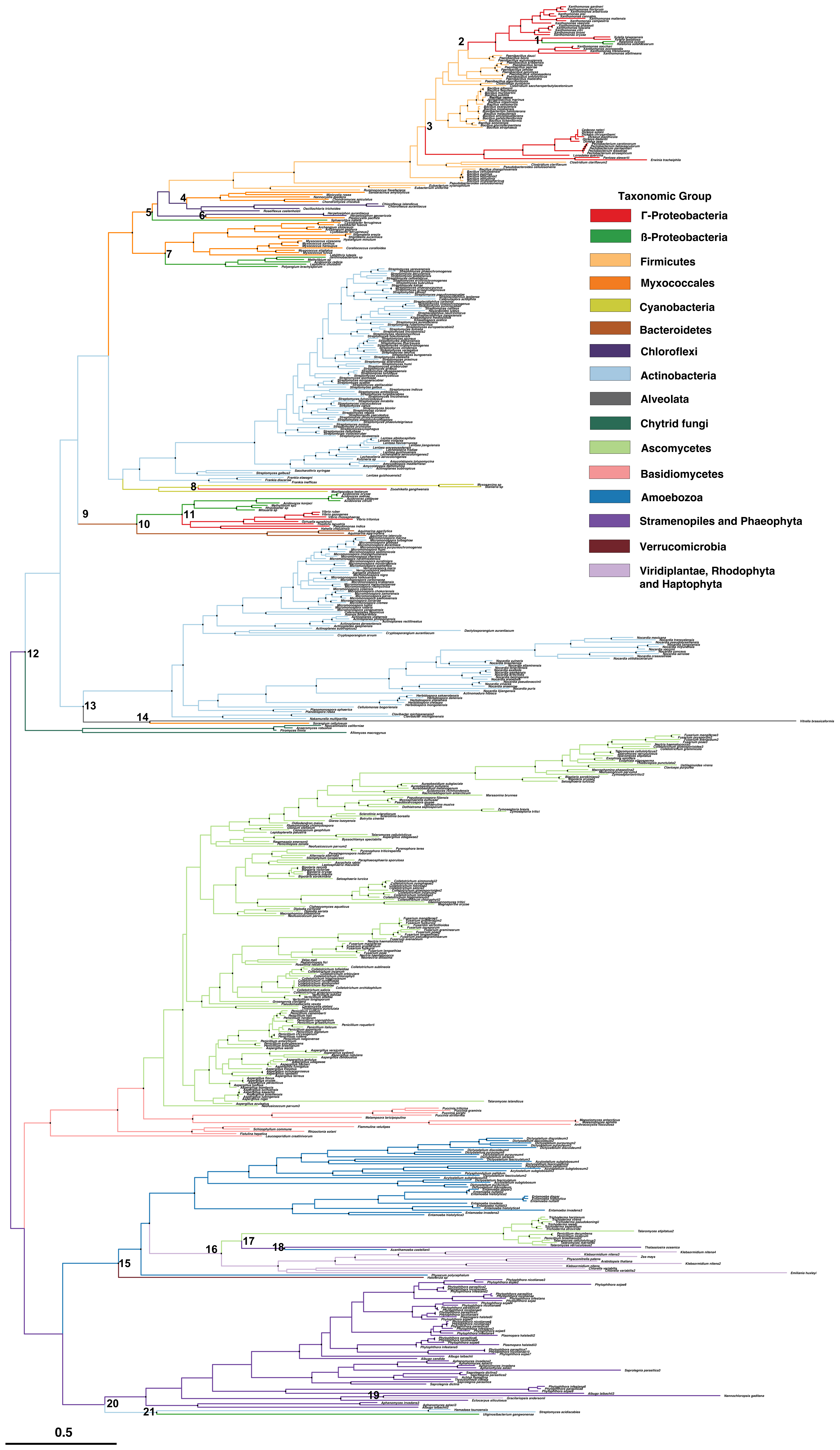

### Supplemental_Figure3

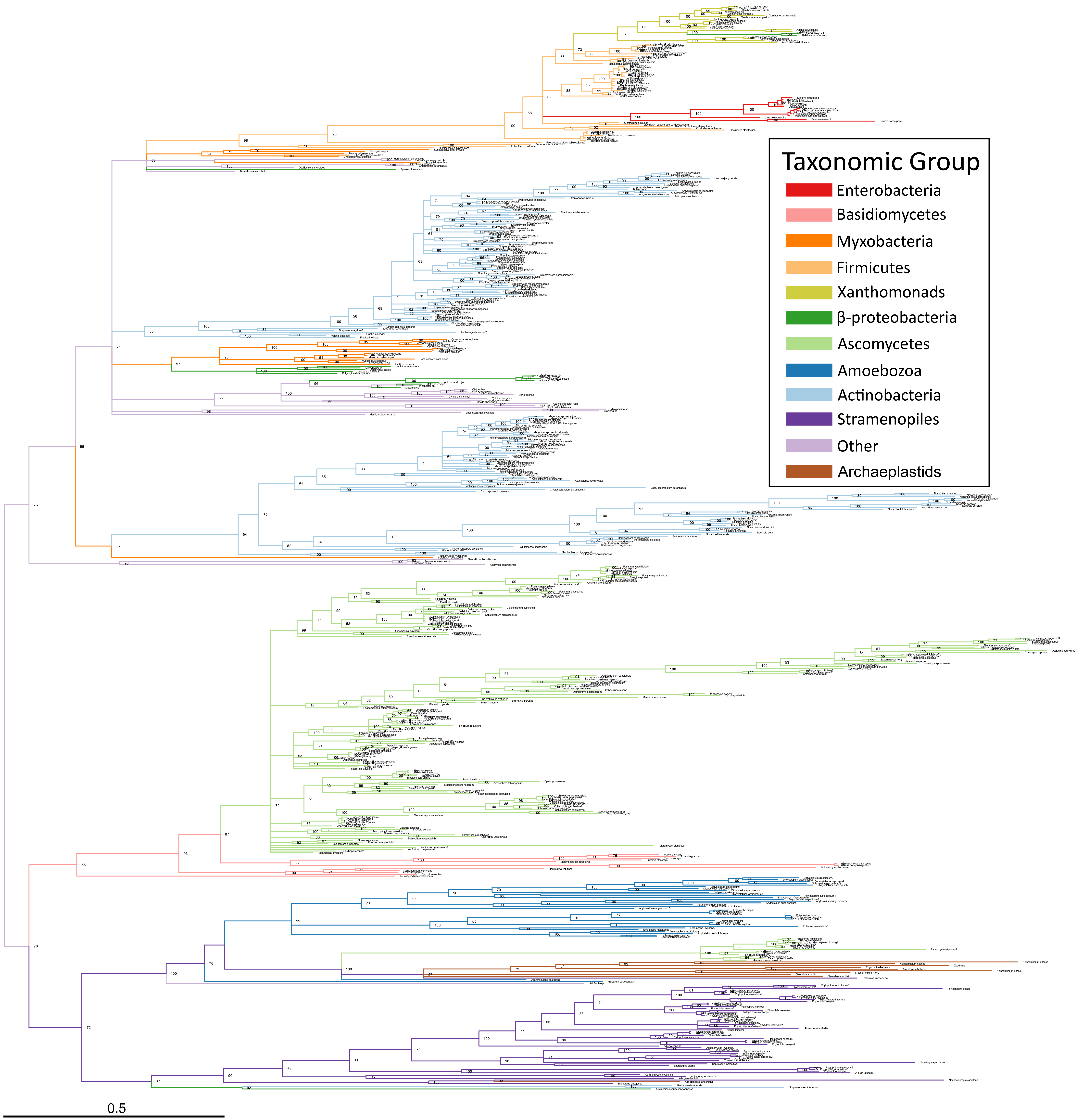

### Supplemental_Figure4

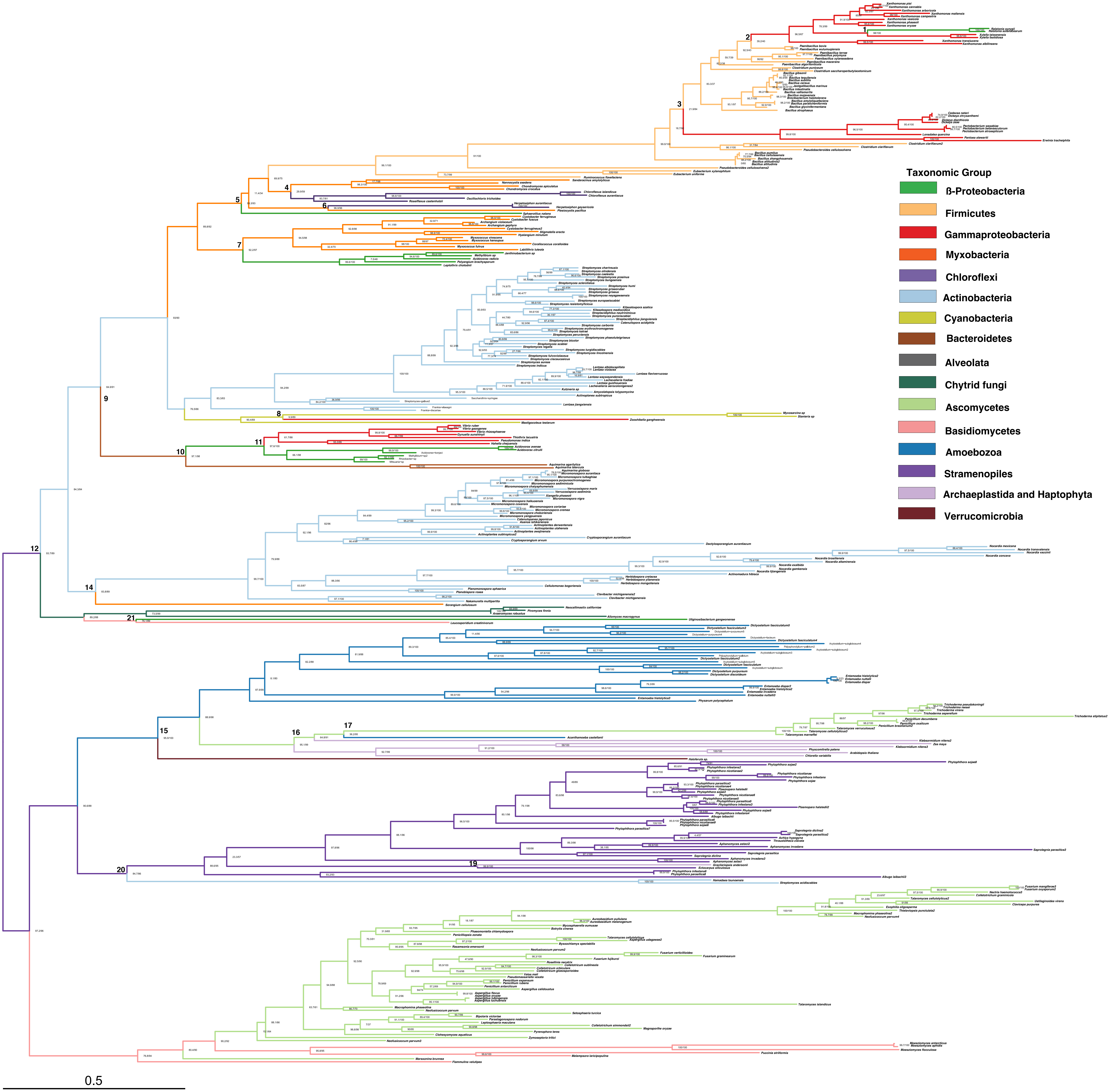

### Supplemental_Figure5

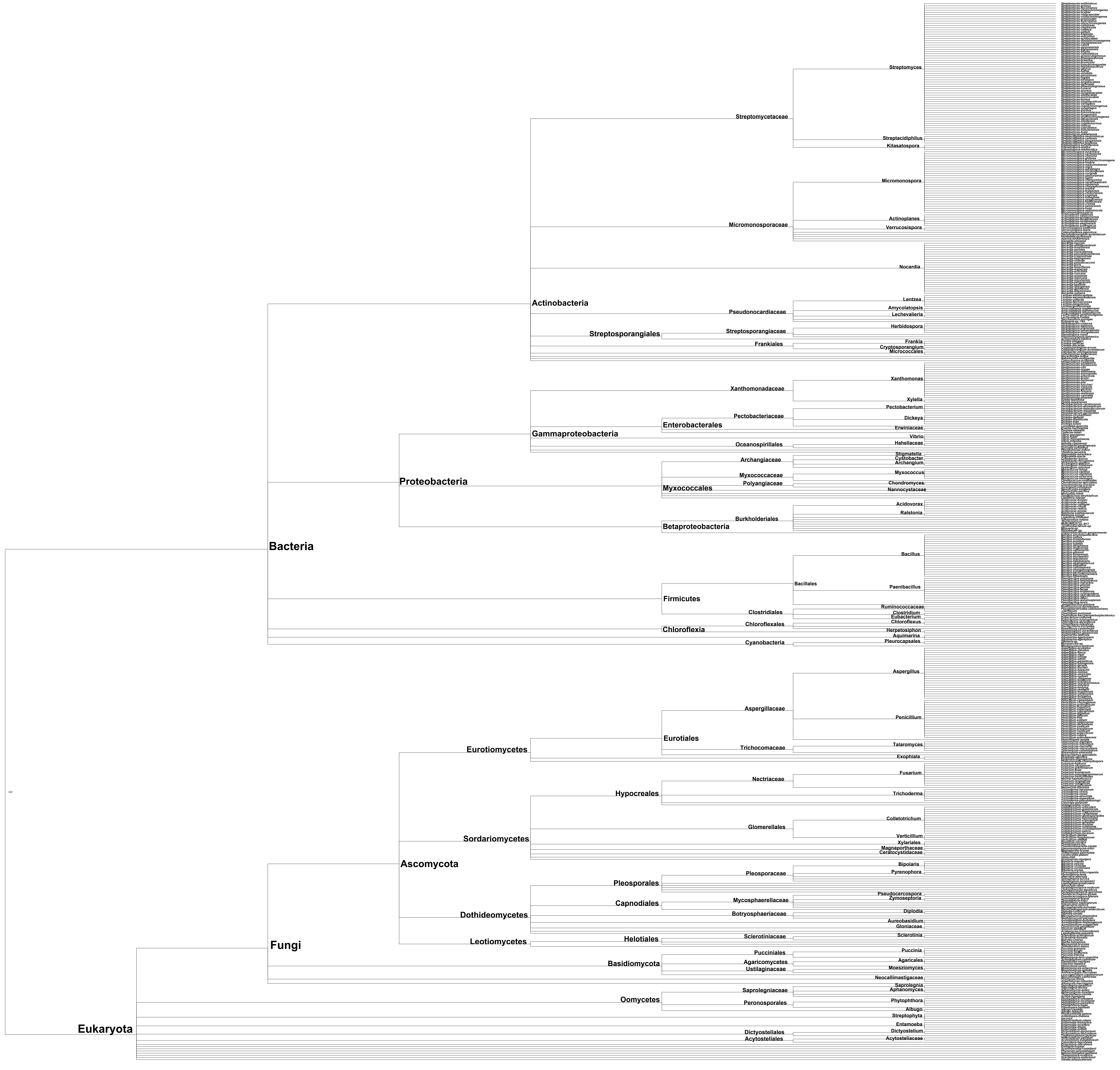

### Supplemental_Figure6

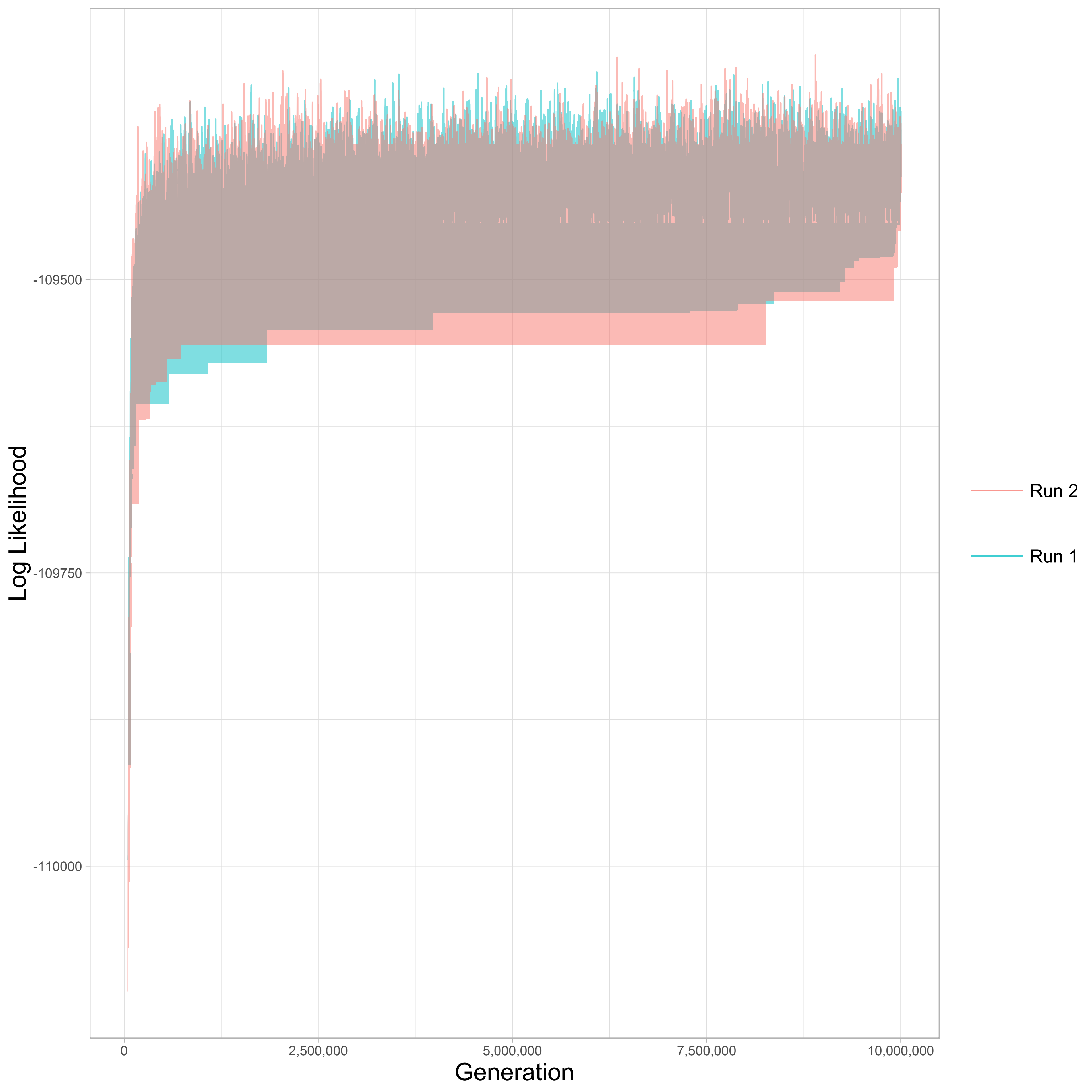
